# Evolution and genetic basis of the plant-penetrating ovipositor, a key adaptation in herbivorous Drosophilidae

**DOI:** 10.1101/2020.05.07.083253

**Authors:** Julianne N. Peláez, Andrew D. Gloss, Julianne F. Ray, Joseph L.M. Charboneau, Kirsten I. Verster, Noah K. Whiteman

**Author notes:** Corresponding authors: Whiteman, N.K.; Pelaez, J.N. These authors contributed equally (Julianne N. Peláez, Andrew D. Gloss).

## Abstract

Herbivorous insects are extraordinarily diverse, yet are found in only one-third of insect orders. This skew may result from barriers to plant colonization, coupled with phylogenetic constraint on plant-colonizing adaptations. Physical barriers have been surmounted through the evolution of key morphological innovations, such as the plant-penetrating ovipositor. Despite their significance, the evolution and genetic basis of such innovations have not been well studied. Ovipositors densely lined with hard bristles have evolved repeatedly in herbivorous lineages within the Drosophilidae. Here, we focus on the evolution of this trait in Scaptomyza, an herbivorous radiation nested in a microbe-feeding clade, sister to Hawaiian Drosophila. Our phylogenetic approach revealed that ovipositor bristle number increased as herbivory evolved. We then dissected the genomic architecture of variation in ovipositor bristle number within S. flava through a genome wide association study. Top associated variants were enriched for transcriptional repressors, and the strongest associations included genes contributing to peripheral nervous system development. Genotyping individual flies replicated the association at a variant upstream of Gαi, a neural development gene, contributing to a gain of 0.58 bristles/major allele. These results suggest that regulatory variation involving conserved developmental genes contributes to a key morphological adaptation required for plant colonization.

## Introduction

Herbivorous insects are among the most successful animal radiations [1, 2], representing approximately one quarter of animal species [3]. Yet, they are found in only one-third of extant insect orders [1], suggesting phylogenetic constraint on adaptations required for this transition. Indeed, the evolution of herbivory requires multi-faceted adaptations, including: locating appropriate host plants, attachment to the host, resisting desiccation, and feeding on nutritionally unbalanced, chemically- and physically-defended plant tissues [4]. Despite the paucity of insect orders with herbivorous species, herbivory evolved many times independently *within* orders [2], including at least 25 times within Diptera [5]. Identifying whether these clades possess key innovations associated with this repeated evolution may help resolve this paradox [6].

The plant-penetrating ovipositor is one such key innovation [7] facilitating entry into this new ecological niche and driving species radiations. It has evolved alongside major species radiations of true fruit flies (Tephritidae), leaf-mining flies (Agromyzidae) and leafhoppers (Cicadellidae), which together comprise ∼27,500 species, as well as lineages of sawflies (Tenthredinidae), katydids (Tettigoniidae), and plant bugs (Miridae) [8]. The insertion of eggs into plant tissue overcomes the challenges of host attachment, desiccation, and access to physically- defended tissues [4]. It also allows neonate larvae to hatch directly into the leaf interior, providing protection from the physical environment and enemies [9]. Some insects with plant-penetrating ovipositors, like agromyzid flies, also consume leaf exudates from oviposition wounds, enabling herbivorous feeding in adults even in the absence of chewing mouthparts, and providing access to novel trophic resources.

The Drosophilidae is a compelling species radiation for studying the plant-penetrating ovipositor as a key innovation for the evolution of herbivory. While most drosophilid species feed on decaying plant tissues, bacteria, and fungi, plant-penetrating ovipositors are found in at least three lineages that evolved herbivory independently: (1) *D. suzukii*, a generalist pest of ripe fruit [10], (2) leaf-mining species within the subgenus *Scaptomyza* (phylogenetically nested within the paraphyletic genus *Drosophila*), which includes the model herbivore *Scaptomyza flava,* a specialized pest of Brassicaceae crops [11], and (3) *Scaptodrosophila notha*, a specialist of living bracken fern (*Pteridium* spp.) [12]. All three lineages bear sclerotized ovipositors, studded with sharp, enlarged marginal bristles used to pierce or scrape into living plant tissue. Drosophilid flies are already models for the evolution of ecological specialization [13] and herbivory, [14] and high-quality genome assemblies across the genus [15] and functional data from *D. melanogaster* enable identification of loci underlying adaptations.

Here, we focus on the evolution of the ovipositor in herbivorous *Scaptomyza,* particularly *S. flava.* In addition to morphological changes to the ovipositor, *S. flava* has acquired a stereotyped behavioral repertoire for leaf puncturing: they tap the ovipositor around the leaf searching for an appropriate location, scoop a hole by repeatedly opening the two oviscapts laterally, then pause briefly, turn counterclockwise, and use their proboscis to imbibe the leaf exudates (Supplemental Videos S1, S2). Females create tens to thousands of punctures per day and oviposit into a small percentage of them [16]. Neonate larvae immediately begin feeding on mesophyll tissue, mining leaves until pupation (Fig. 1a). Those hatching outside of the leaf do not survive [11].

**Figure 1.**
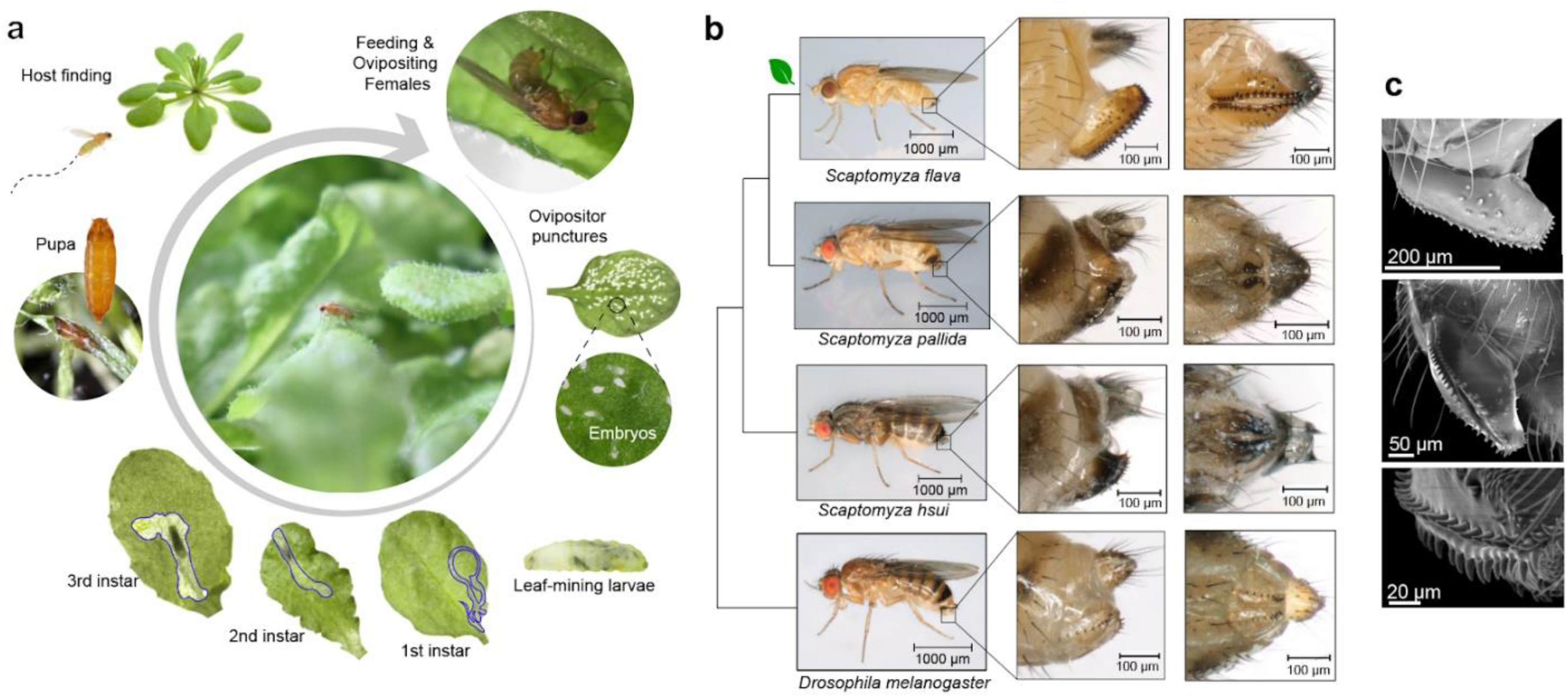
The morphology of the female ovipositor of the herbivorous drosophilid *Scaptomyza flava* enables cutting into tough plant tissues. (a) The life cycle of *S. flava* is strongly dependent on host plants for female nutrition and larval development. On the underside of an *Arabidopsis thaliana* leaf, a female uses her serrated ovipositor to scoop a leaf puncture for feeding and egg- laying. Larval mines outlined in blue. (b) Comparison of the ovipositors (insets) of herbivorous and non-herbivorous drosophilid species. (c) Scanning electron micrographs of the ovipositor of *S. flava*.

Although the ovipositors of herbivorous drosophilids differ noticeably in many aspects of their shape and size, most species (exceptions include *Scaptomyza flavella* [17] and *Scaptodrosophila megagenys* [12]) share a single row of supernumerary bristles along the ventral margin (e.g. Fig. 1b, c). We therefore focused on bristle number, which has been well-studied from a quantitative genetics and developmental biology perspective [18]. We first investigated whether ancestral increases in ovipositor bristle number paralleled the transition to herbivory in *Scaptomyza*, using phylogenetic generalized least squares (PGLS) methods and ancestral state reconstruction (ASR). To understand the nature of mutations (i.e. coding versus regulatory, mono- versus polygenic, and candidate developmental processes involved) that could have given rise to increased ovipositor bristle number, we used pooled genome-wide association mapping (pool-GWAS) [19, 20] within the herbivorous species *S. flava*. Finally, we sought to validate our pool-GWAS by genotyping individuals and estimating the effect size of a single nucleotide polymorphism (SNP) that reached genome-wide significance.

## Materials and Methods

### (a) Phylogeny reconstruction

We estimated a phylogeny of *Scaptomyza,* including Hawaiian *Drosophila*, the sister clade of *Scaptomyza*, using 11 genes in 95 taxa (Table S1). We expanded a previous dataset [21] with five additional taxa: two with sequenced genetic markers, *S.* nr. *nigrita* (Nevada) and *S. montana* (Arizona) [22], and three obtained in this study from California, *S.* nr. *nigrita*, *S. montana*, and an undescribed species. DNA extraction and PCR methods were described previously [23]. PCR amplicons were cleaned and Sanger sequenced in both directions at the UC Berkeley Sequencing Facility, trimmed and manually aligned to the other taxa [21] in Geneious v.10.0.5. We estimated a species tree by maximum likelihood (ML) in RAxML [24], and a time-calibrated species tree by Bayesian inference using MrBayes v.3.2.4 [25] and BEAST v.2.4.6 [26]. Alignment partitioning and model implementation are described in the Supplementary Methods. Complete phylogenies are reported in Fig. S1 and S2.

### (b) Ovipositor trait evolution

To test whether ovipositor bristle number changed significantly during the evolution of herbivory, we performed PGLS [27] on ovipositor bristle number, accounting for larval diet [23] and phylogenetic relatedness. We collected bristle counts from illustrations or images from the literature, or from wild or lab-reared individuals (Table S2). Because distinguishing between bristle types was not always clear, we counted all bristles, regardless of morphology, position, or size, on an oviscapt, averaging across multiple literature sources if available. Ovipositors of wild and lab-reared flies (n ≤ 10 per species) were mounted onto slides with Permount mounting medium (Fisher Scientific) and coverslips. Ovipositors were photographed using an EOS Rebel T3i camera (Canon) mounted on a Stemi 508 stereo microscope (Zeiss) with a 1000 μm scale bar.

PGLS regression was performed using ape [28] and picante [29] packages in R. Models of trait evolution (Brownian motion, Ornstein–Uhlenbeck, Early Burst, and white noise) for bristle number were compared using AICc in the geiger R package [30], and Brownian motion was selected as the best fit (Table S3). The degree of phylogenetic signal in the residuals was estimated using Pagel’s lambda (λ) [31]. To visualize correlated evolutionary changes in diet and bristle number, ASR of both traits were estimated by ML using phytools [32] and ape [28] and mapped onto the phylogeny. Models of trait evolution (equal rates, symmetric, and all rates different) for larval diet were compared, and equal rates was selected as the best fit (Table S4).

To investigate whether ovipositor length and/or body size influences ovipositor bristle number, bristle number was linearly regressed against ovipositor and thorax length (proxy for body size). We used an expanded set of taxa included in a published dataset from [33] where all three measurements were taken consistently across 67 species, supplemented with direct measurements from four additional species (Table S5). Ovipositor bristle counts were averaged across oviscapts and across individuals within a species. Individuals were excluded if only a subset of bristles were counted.

### (c) Mapping population and measurements for pool-GWAS

To identify specific genetic polymorphisms contributing to variation in bristle number, we used a pool-GWAS to detect allele frequency differences between pools of individuals with extreme phenotypes from the same population. Two *S. flava* outbred laboratory populations were founded from larvae collected from mustard plants, one individual per plant, near Dover, New Hampshire, USA: 79 larvae from *Turritis glabra* (referred to as “NH1”) and 58 from *T. glabra* and *Barbarea vulgaris* (“NH2”). After eclosion, adults were transferred to one mesh cage per population, containing *Arabidopsis thaliana* (Col-0 accession). Over 200 F1 offspring per population were reared on a mixture of *T. glabra* and *B. vulgaris*, and adult female F2 offspring were preserved in 95% ethanol and phenotyped for GWAS.

Ovipositors were mounted on slides as described above. Bristles were counted along the ventral edge (Fig. 3a), excluding largely invariable apical bristles. We quantified ovipositor length and wing chord (proxy for body size) using ImageJ. Wing chord was measured from the base to the wing apex following the third longitudinal vein (Fig. 3a). Two independent measurements were averaged per specimen. Linear regression analyses in a pilot experiment (N = 100, NH1 and NH2 flies) revealed that bristle number was positively correlated with ovipositor length (*B* = 0.097 [S.E. = 0.025] pegs per μm length, *R^2^* = 0.134, *P* = 0.0001), but not wing length (*B =* 0.001 [S.E. = 0.002], *P* = 0.25). We therefore quantified both ovipositor length and bristle number for all individuals (NH1, N = 308 flies; NH2, N = 422 flies).

Narrow-sense heritability of ovipositor length and bristle number were quantified using mother-daughter regression; further details are presented in the Supplemental Methods.

### (d) Pooled genome sequencing

Flies in the NH1 and NH2 populations were split into two phenotypically extreme pools per population (four pools: NH1-low, NH2-low, NH1-high, NH2-high), composed of 60-85 females in either the upper or lower 20% tail of the distribution of residual bristle number. Residual bristle number was determined through a linear regression of ovipositor bristle number against ovipositor length using the *lm* function in R. Flies were homogenized with a stainless-steel bead and TissueLyser (Qiagen). Genomic DNA was extracted using a DNeasy Blood and Tissue Kit (Qiagen). One Illumina library per pool was constructed with 100 bp paired-end reads and a 350 bp insert size, and each library was sequenced on one half lane on an Illumina HiSeq 2500 at Arizona State University.

### (e) Read mapping, pool-GWAS and gene ontology enrichment analysis

Reads were mapped to the *S. flava* reference genome (GenBank accession no. GCA_003952975.1) and filtered following best practices for pooled genome sequencing [34]. Statistical significance of between-pool allele frequency differences per site was estimated using the Cochran-Mantel-Haenszel test [35]. See Supplemental Methods for further details. To identify genes located in or near the top SNPs, ranked by *p* value, we viewed the *S. flava* genome assembly and gene annotations [11] in Geneious v.10.2.6 and located the nearest annotated gene in either direction from the SNP. We checked for unannotated genes between the SNP and closest annotated gene by comparing the spanning sequence against the *D. melanogaster* RefSeq protein database, using NCBI BLASTx with default settings. Information on gene function was collected from the Gene Summary, Gene Ontology Annotations, and linked publications in Flybase (release 2020_01) [36]. To aid in interpretation of the pool-GWAS results, we profiled linkage disequilibrium (LD) in a wild population of *S. flava* near the NH1/NH2 collection locality. Further details are presented in the Supplementary Methods.

To determine if any predicted functions were overrepresented among genes intersecting the top associations, we performed a Gene Ontology enrichment test using GOWINDA, which implements a permutation-based approach tailored to the properties of GWAS datasets [37]. Full details, including orthology-based functional annotation and extension of gene models to capture regulatory regions, are described in the Supplemental Methods.

### (f) Replicating pool-GWAS association for a candidate SNP

Pool-GWAS can be confounded by uneven contributions of individuals to pools and biases in sequencing and read mapping. To replicate our pool-GWAS results using an approach robust to these confounding factors, we genotyped individuals at one of the top SNPs and estimated its effect size (Fig. 3g; Table S6). The SNP was chosen because of its position upstream of *G alpha i subunit* (*Gαi*), a gene involved in asymmetric cell division of sensory organ precursor (SOP) cells from which bristles are derived [38]. Ovipositor bristle number and length were measured as described above. Genomic DNA was extracted from 74 females from NH1 and NH2 mapping populations, and a target region of 500bp around the SNP was Sanger-sequenced. Additional details are presented in the Supplementary Methods.

Bristle number was modeled in a generalized linear model, assuming an additive effect of the major allele, using the *lm* function in R. The model accounted for collection locality, lab-rearing host plant species, ovipositor length, and whether they were included in the high or low bristle number pools (to account for polygenic effects of the genomic background). Effect size (β) was estimated as the shift in bristle number (in standard deviations) expected from a single allelic substitution. A second model tested the effects of these factors on ovipositor length.

Pairwise measures of LD between the SNP and other variant sites (minimum frequency >0.05) were calculated using the *LD2* function from the pegas package in R [28]. Correlation among alleles is given by *δ* [39] with strong LD indicated by | *δ* | > 0.5 (*p* value < 0.01).

## Results

### (a) The evolution of herbivory coincided with an increase in ovipositor bristle number

PGLS methods revealed that ovipositor bristle number is strongly influenced by larval diet (*F*1, 1=4.33, *P* = 0.05) and phylogenetic relatedness (Pagel’s λ = 1) (Table S7). Ancestral state reconstructions of bristle number and larval diet similarly suggest that ovipositor bristle number increased coincident with the evolution of herbivory in *Scaptomyza*, estimated ∼10.4 million years ago (mya) (8.2 -13 mya, 95% highest probability density) (Fig. 2a; Fig. S3). Relative to interspecific differences, variation within species is low (Fig. 2b).

**Figure 2.**
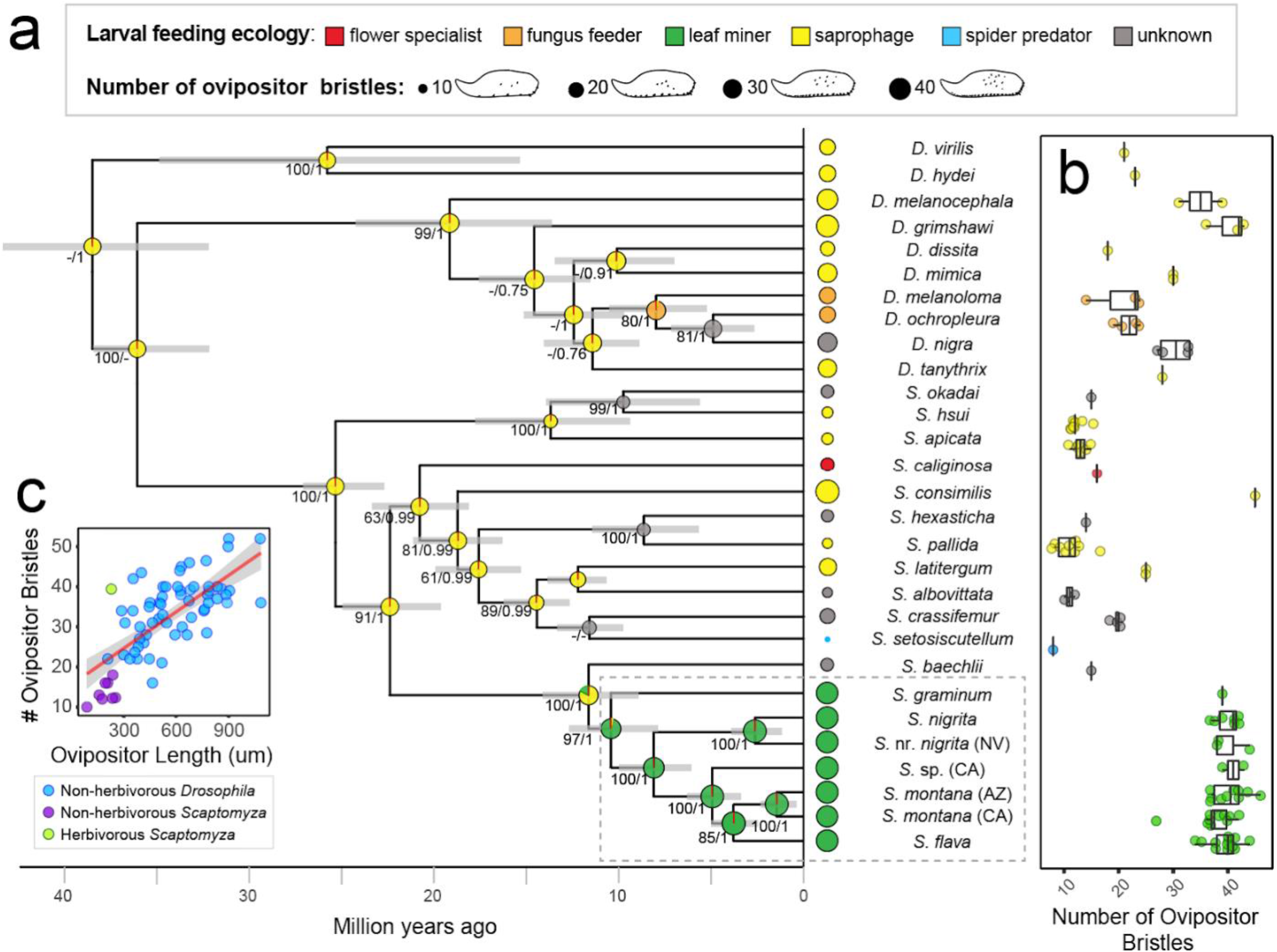
The evolution of herbivory within *Scaptomyza* coincides with an increase in ovipositor bristle number. (a) Time-calibrated phylogeny of herbivorous *Scaptomyza* and their non-herbivorous relatives, based on ML and Bayesian analyses, using 11 genes and fossil and biogeographic time calibrations. Branch support is indicated by ML bootstrap values (≥50%) and Bayesian posterior probability (≥0.9). Bars at nodes indicate 95% highest posterior density interval around the mean node age. Pie graphs at nodes show probabilities of ancestral larval diets, and size represents ancestral ovipositor bristle number (per oviscapt) estimated from ML ASR. Average bristle number for extant species are shown at the tips, with individual counts shown in (b). (c) Scatterplot of ovipositor bristle number as a function of ovipositor length.

The increase in ovipositor bristle number in herbivorous *Scaptomyza* is likely to reflect increased bristle density, rather than increased ovipositor or body size. Regressing bristle number onto ovipositor length and female thorax length (a proxy for body size), we found that while bristle number is strongly predicted by ovipositor length across drosophilid species (β = 0.53, t(53) = 4.3, *P* < 0.001, adjusted R^2^ = 0.47; Fig. 2c, Table S8), the herbivorous species *S. flava* has proportionally more ovipositor bristles relative to ovipositor length than non-herbivorous species (highest residual value; Fig. S4). Further, ovipositor bristle number was not strongly predicted by thorax length (β = 0.24, t(53) = 1.9, *P* = 0.06).

### (b) GWAS on ventral ovipositor bristle number

Variation in ovipositor bristle number was normally distributed in the NH1 and NH2 outbred laboratory populations of *S. flava* (Fig. 3a-b), typical of a quantitative trait controlled by multiple loci. Linear regression of ovipositor bristle number from mother-daughter pairs, controlling for the effect of ovipositor length, revealed that additive genetic variation accounted for roughly half of this phenotypic variation (*P* = 0.034, *h^2^* = 0.50 ± 0.27 SE; Fig. 3c). By contrast, variation in ovipositor length was not heritable (*P* = 0.31).

**Figure 3.**
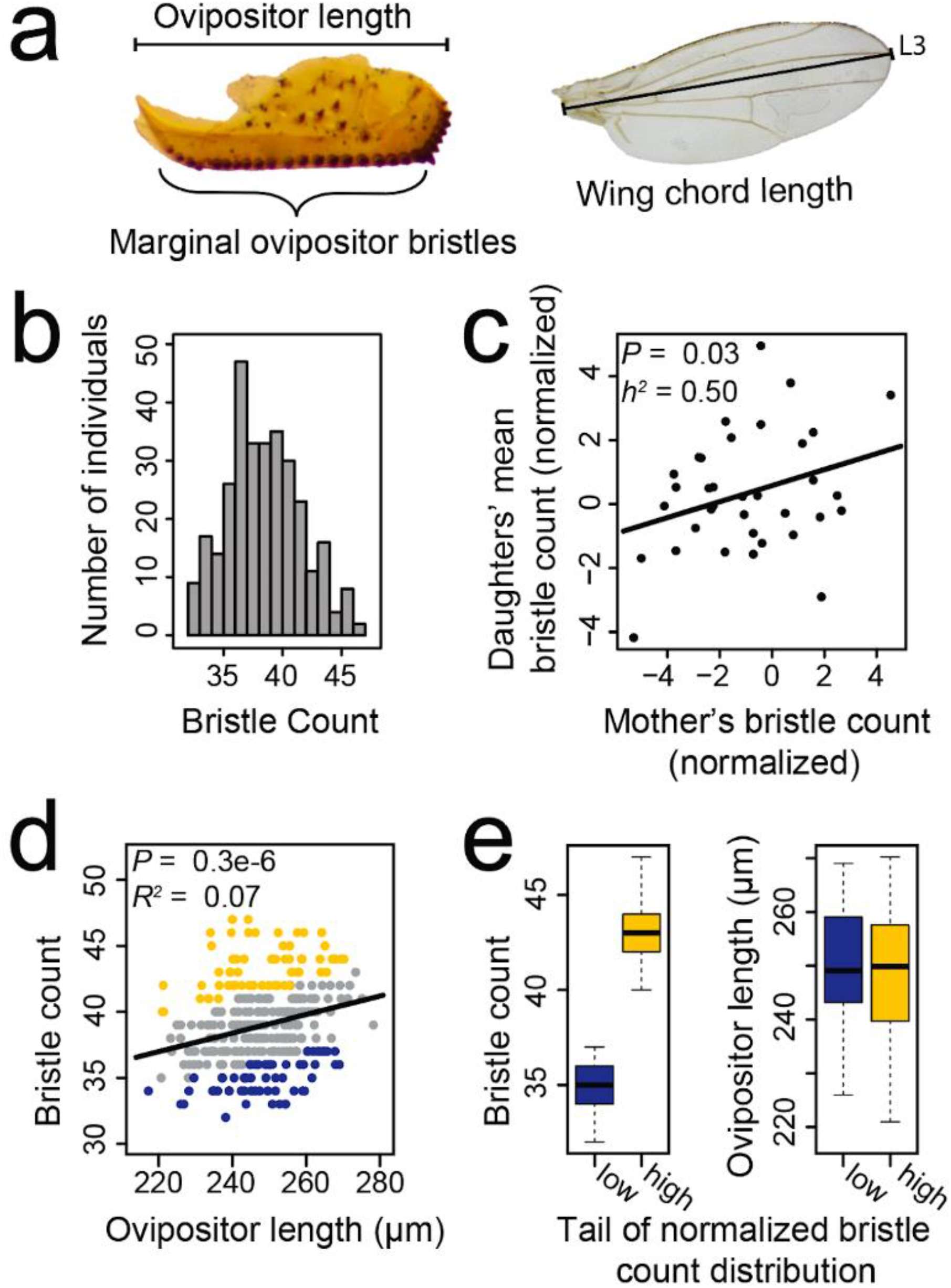
Variation in the number of plant-cutting ovipositor bristles is normally distributed and heritable in *S. flava*, enabling quantitative genetic dissection. (a) Ovipositor bristle counts include those lining the ventral margin, summed across both oviscapts. Wing chord length was measured along the third longitudinal vein (L3). Panels b, d, and e show phenotype distributions used for the pool-GWAS in the NH1 outbred mapping population. **(b)** Ovipositor bristle number follows a normal distribution. (c) Ovipositor bristle count, expressed as residuals from a linear regression of bristle count against ovipositor length, is heritable in the narrow sense (*h^2^* = 0.50) from mother-daughter regression analysis (N = 35). **(d)** After regressing out the effect of ovipositor length on bristle count, pools of phenotypically extreme individuals were constructed for genome sequencing by combining individuals in the upper (yellow) or lower (blue) 20% tails of the distribution. **(e)** Individuals in the low pool had ∼20% fewer bristles, but not statistically different ovipositor lengths, than those in the high pool.

We sought to characterize the genomic architecture underlying this variation using a pool-GWAS. Because ovipositor length was correlated with bristle number (Fig. 3d), low and high bristle number pools were constructed with bristle number adjusted relative to that expected from ovipositor length (Fig. 3e). Our pool-GWAS approach should therefore interrogate bristle number independently of ovipositor size, while also minimizing noise introduced by non-heritable variation in ovipositor length that could otherwise impede GWAS. Whole genome re-sequencing of the four pools were mapped to the *S. flava* genome, resulting in a mean experiment-wide coverage depth of 166X per polymorphic site. After excluding low frequency variants (1.6 million SNPs remaining), we found an excess of SNPs with significantly differentiated allele frequencies among high and low bristle number pools (Fig. 4a), with 5 and 19 significant SNPs at 5% and 10% false discovery rate (FDR) cutoffs, respectively (Table 1; Table S9). Because LD decays in *S. flava* at a rapid rate similar to that seen in *D. melanogaster* (Fig. 4b), SNPs showing the strongest associations are likely in close proximity to causal polymorphisms or are causal themselves.

**Table 1.** Top SNPs associated with variation in ovipositor bristle number are located in or near genes involved in the development of bristles, cuticles, and the nervous system. SNPs reaching genome-wide significance (FDR⩽0.05) from the pool-GWAS are shown in descending *P*-value ranking.

| P-value ranking | Scaffold [scaffold length] | Position on scaffold | P-value | FDR q value | Nearby gene(s) | SNP location relative to gene | Gene function |
| --- | --- | --- | --- | --- | --- | --- | --- |
| 1 | scaffold00465 [104,011] | 184 | 4.66E-09 | 0.008 | **Muscle-specific protein** | 453 bp downstream | Actin binding; cytoskeleton organization; required for proper positioning of muscle nuclei, mitochondria, and neuromuscular junction |
| 2 | scaffold00015 [769,991] | 186,025 | 4.3E-08 | 0.036 | **heavyweight** | 8,623 bp downstream | Predicted to have phosphotyrosine residue binding activity; polymorphisms associated with body mass and starvation resistance |
|  |  |  |  |  | **cuticular protein 11B** | 1,625 bp downstream | Chitin-based cuticle development |
| 3 | scaffold00063 [441,164] | 140,206 | 6.69E-08 | 0.037 | **G protein alpha i subunit** | 2 bp upstream | Asymmetric neuroblast division and asymmetric protein localization involved in cell fate determination; cytoskeleton organization; and nervous system development |
| 4 | scaffold00053 [434,434] | 135,098 | 1.20E-07 | 0.048 | **sloppy paired 2** | 560 bp upstream | Transcription factor that regulates embryonic segment polarity and neural fate specification by temporal patterning of medulla neuroblasts |
|  |  |  |  |  | **CG11018** | 1,809 bp upstream | Unknown |
| 5 | scaffold00071 [361,359] | 290,082 | 1.42E-07 | 0.048 | **CG32655** | 9,932 bp downstream | Unknown |
|  |  |  |  |  | **tenascin accessory** | 42,315 bp downstream | Nervous system development; regulation of cell-cell adhesion; and synapse organization |

Note: SNPs with exceptionally high coverage (likely erroneous; 2 among top 25) were filtered.

**Figure 4.**
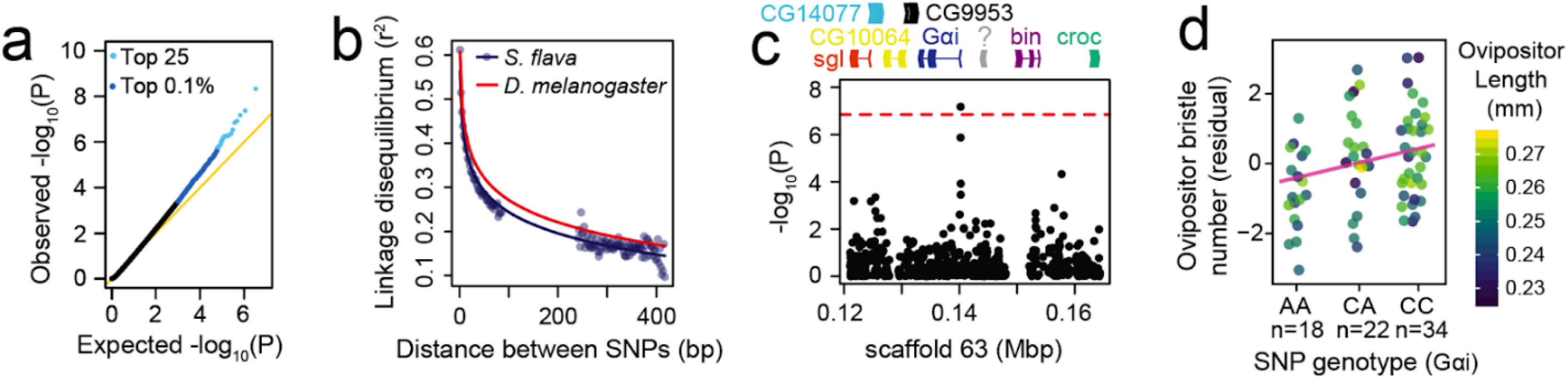
Pool-GWAS for variation in *S. flava* ovipositor bristle number implicates genes involved in nervous system development. (a) An excess of strong *P*-values suggests an enrichment of true associations among the top scoring SNPs. **(b)** The relationship between physical distance and linkage disequilibrium, inferred from pooled sequencing of wild *S. flava*, is similar to that seen in *D. melanogaster*. **(c)** Manhattan plot centered on a top SNP upstream of *G-alpha i subunit* (*Gαi*), a gene that functions in nervous system development [38]. The red line indicates the 5% FDR cutoff for genome-wide significance. Annotated genes are plotted above; ambiguous orthology indicated by “?”. **(d)** Genotyping individuals for the SNP near *Gαi* replicates the pool-GWAS result. Bristle number, expressed as residuals generated by subtracting predicted values based on covariates from observed values, increases additively with each major allele and independently of ovipositor length, shown by color scale. Regression line shown in pink.

Many of the top SNPs (Table S9), including those reaching genome-wide significance (FDR⩽0.05, Table 1), were located near genes involved in neural development or neural cell fate specification (i.e. *G protein alpha i subunit, sloppy paired 2, tenascin accessory*), cytoskeleton organization (i.e. *muscle-specific protein*), and cuticle development (i.e. *cuticular protein 11B*).

### (c) Gene ontology enrichment analysis on candidate SNPs

To gain insight into developmental and physiological mechanisms that may contribute to variation in ovipositor bristle number, we tested for enriched gene ontology (GO) annotations among genes intersecting SNPs with the strongest pool-GWAS associations (top 0.1% and 0.005% of *P*-values genome-wide). Using a restricted set of GO terms to minimize redundancy, we uncovered a single enriched term: RNA polymerase II-specific DNA-binding transcription repressor activity (GO:0001227; Table 2). Transcriptional repressors fine-tune gene expression levels during the specification of cell fate during development [40]. Notably, the strongest pool-GWAS association among transcriptional repressors falls in the *S. flava* gene orthologous to *hairy* (*h*) in *D. melanogaster* (Table S10), which functions in the establishment of bristle precursor positioning from within proneural clusters [41].

**Table 2.**
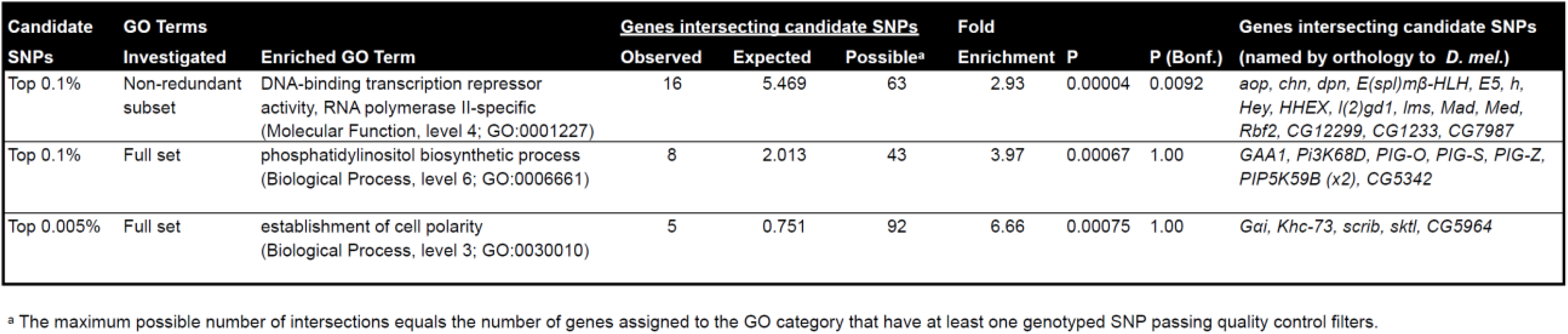
Gene ontology terms enriched among genes intersecting the most significant pool-GWAS SNPs.

We further tested for enrichment using the exhaustive list of all GO terms. This approach imposes a conservative multiple testing burden, and no terms were enriched after applying a strict Bonferroni correction. However, two terms surpassed a nominal cutoff of *P* < 0.001, and both reflect broadly conserved developmental functions in eukaryotes: phosphatidylinositol (PI) biosynthetic process and establishment of cell polarity (Table 2). Many of the candidate genes annotated with PI biosynthetic process (GO:0006661) are kinases and transferases involved in production of PI derivatives (Table S10), which act as signaling molecules that regulate cellular growth and patterning [42–44]. Notably, establishment of cell polarity (GO:0030010) precedes the differentiation of sensory organ precursors into distinct neural cell types through asymmetric cell division [45]. *G protein α i subunit* (*Gαi*), one such gene involved in polarization and asymmetric division of neural cells [38], harbored one of the strongest pool-GWAS associations in our study, surpassing the 5% FDR threshold for genome-wide significance (Fig. 4c, Tables 1 and S8).

### (d) Replication of a top candidate SNP from pool-GWAS

To validate the pool-GWAS, we focused on a SNP in the 5’ UTR of *Gαi*, one of the strongest pool-GWAS associations. We phenotyped and genotyped individual adult females and recapitulated the pool-GWAS findings. Bristle number increased by 0.58 per major allele carried (*β* = 0.11 standard deviations, t(68) = 2.88, *P* < 0.005; Fig. 4d; Table S11). This SNP explained 9.5% of the total variance in bristle number (partial adjusted *r*^2^). As expected given our study design, the SNP did not have an effect on ovipositor length (*β* = 0.02 standard deviations, t(69) = 0.177, *P* > 0.05; Table S12). Out of 5 variant sites (≥0.05 min. freq.) in the sequenced region, two were in strong LD with the focal SNP and were located upstream of *Gαi*’s coding sequence or in an intronic region (Table S13). Further study will be necessary to identify the causal variant(s) in this region.

## Discussion

The plant-penetrating ovipositor of herbivorous insects presents an excellent opportunity to study the evolution and genomic architecture of a complex trait, given its clear adaptive role in egg-laying and the quantitative nature of ovipositor morphological traits. We focused on the evolution of the ovipositor in the genus *Scaptomyza,* in which herbivory has evolved relatively recently, ca. 10.4 mya. The wealth of data from the *Drosophila* literature made our analyses possible: genitalic data from numerous taxa to investigate macroevolutionary shifts in bristle number, and knowledge of the genetics and development of bristle number in *D. melanogaster* to understand the genetic architecture underlying variation at the population level in *S. flava*.

From a macroevolutionary perspective, we found that ovipositor bristle number underwent a marked increase that coincided with the evolution of herbivory within *Scaptomyza*, a significantly larger shift than expected from the distribution of background rates of evolution across the phylogeny (Fig. 2a). Surprisingly, we also found that ovipositor bristle number is an evolutionarily malleable trait, repeatedly increasing and decreasing across the phylogeny, with a five-fold range across *Scaptomyza*. High variability was similarly seen *within* species, with a 1.5- fold range in *S. flava*. The lack of strong evolutionary constraint over both macro- and microevolutionary timescales, along with availability of heritable standing genetic variation within populations, suggests that increased ovipositor bristle number is highly accessible to adaptive evolution. If these patterns hold in other Drosophilidae, the evolutionary malleability and accessibility of this trait may help explain why densely bristled ovipositors have convergently evolved across independent transitions to herbivory, such as the lineages that include *D. suzukii* and *S. notha*.

While the evolution of increased ovipositor length has been studied in *D. suzukii* as a key trait to facilitate cutting into ripe fruit [46], our phylogenetic analyses revealed that bristle number was still highest in herbivores even after accounting for ovipositor length (Fig. 2c, Fig. S4), suggesting that bristle number increased not simply as a result of ovipositor elongation, but from increased bristle density. Narrow sense heritability estimates of bristle number (adjusted for ovipositor length) in *S. flava* further showed that bristle number was heritable (Fig. 3c), while ovipositor length was not. Using ovipositor length-adjusted bristle counts, our GWAS thus targeted variation in bristle number and identified associated genetic variants contributing to this aspect of *S. flava*’s ovipositor.

Pinpointing the genetic changes that gave rise to adaptive traits that evolved millions of years ago can be difficult because genetic architectures may differ over short versus long timescales [47]. Still, GWAS can illuminate genes and gene functions that shape standing phenotypic variation and may contribute to evolution over longer timescales. Our GWAS on ovipositor bristle number indicates that broadly conserved developmental genes and processes play a role in ovipositor bristle density. Genes encoding transcription repressor proteins were enriched near the strongest GWAS associations, and many top-scoring SNPs were located near genes with known roles in neural development, including *Gαi* [38] and *slp2* [48]. This is consistent with our understanding that insect bristles are developmentally derived from single neural precursor cells (sensory organ precursors or SOPs) that differentiate through asymmetric cell divisions to generate mechanosensory and chemosensory neurons that innervate bristles and the cells that form their shaft, socket and sheath [49]. Innervation of ovipositor bristles has been demonstrated in flies, including in *D. melanogaster* [50]. Tinkering with genes involved in neural or SOP development could presumably lead to increased cell divisions specific to these SOP lineages to produce more bristles. In other *Drosophila* species, genes involved in neural development underlie differences in bristle number on male genitalia, sexcombs of the forelegs [51] and the thorax [52]. Specifically, *hairy* (*h*) was the top-scoring SNP within the GO category most over-represented in our strongest GWAS hits. This gene has been implicated in both within-species and between-species variation for several bristle traits in *D. melanogaster* and its close relatives [53, 54]. RNAi knockdown of *h* in *Drosophila* has validated its involvement in male genital development, specifically clasper size and bristle number [54]. Intriguingly, *h* falls within a narrowly-mapped genomic region underpinning divergence in clasper bristle number among sister species of *Drosophila* [54]. Its role in bristle and genital development, along with its contribution to intra- and inter-species variation in bristle number, make *h* an excellent candidate for ovipositor bristle variation. It also highlights intriguing potential for genetic parallelism for variation in bristle number across the body, between sexes and across species.

Studies on the genetic architecture of adaptive traits have largely focused on monogenic, Mendelian traits with large effect sizes of candidate loci [55–57] with lower detection thresholds than genetically complex traits. Ovipositor bristle number represents a tractable quantitative trait for genetic dissection because of its meristic nature, high variability and heritability. Despite having a genetic basis similar to many quantitative traits — many small effect SNPs underlying variation — we still were able to detect a SNP with moderately large effect (validated by individual genotyping). Our results suggest that pool-GWAS can be a viable method for pinpointing genomic regions that underlie quantitative trait variation. Candidate SNPs can then be interrogated through functional experimentation to understand how alternative alleles influence cell division, size expansion, and reorganization during development [46]. Focusing on the developmental pathways, genes, and regulatory regions identified through our GWA mapping would offer a future route to illuminate how incremental changes could have created this key innovation in herbivorous insects.

## Data accessibility

All data files and scripts were deposited in the Dryad Repository (https://doi.org/10.6078/D1841H). Sanger sequences for estimating the *Scaptomyza* phylogeny were uploaded to GenBank (MH938262-MH938270). Available at NCBI sequence read archive are Illumina sequences for the pool-GWAS (SRR11252387-SRR11252390), and to evaluate linkage disequilibrium (SRR15275350-SRR15275365). Sanger sequences for replicating the *Gai* SNP effect size were deposited on GenBank (MH884655-MH884734).

## Acknowledgements

We thank Mitchell Feldmann and Amelia White for caring for fly populations, Aaron Pomerantz for assistance imaging ovipositors, and Justin Lack for sharing advice and scripts for read mapping. We also thank Craig Miller, Kristin Scott, Michael Nachman, and Rasmus Nielsen for their feedback and advice.

## Supplemental Methods

### Alignment partitioning and model implementation for phylogeny reconstruction

The concatenated alignment was partitioned by codon and gene, with ribosomal genes given single partitions. The ML analysis used a GTR+gamma model on all partitions. To evaluate consistency across runs, five independent runs were performed, with distinct starting seed. Each run included 1,000 bootstrap replicates, and a slow ML search on every 5th tree. The phylogeny with the highest likelihood was used for ancestral character estimation. For the Bayesian analysis, models of sequence evolution were selected for each partition with the Akaike information criterion (AIC), using MrModeltest2 v.2.3 [1] and PAUP* v.4.0a [2]. To infer a phylogeny and divergence times, a Markov-Chain Monte-Carlo (MCMC) analysis was performed as previously described using the same time calibrations points and run parameters [3]. To ensure that the Markov chain adequately converged to a stationary distribution, forty replicate runs of 10 million generations each were performed and implemented in BEAST v.2.4.6 [4] with BEAGLE [5] for multicore processing. The first ten percent of samples were discarded as burn-in. Trees were re-sampled every 250,000 generations, and combined using LogCombiner v.2.4.6 [4]. Tracer v.1.6.0 [6] was used to confirm that ESS values were sufficient for reliable parameter estimates (ESS >200).

### Narrow-sense heritability estimates from mother-daughter regression

Fifty single male-female pairs (virgin females) from the combined NH1 and NH2 colonies were individually mated on *T. glabra* in Magenta boxes (Sigma-Aldrich) with mesh covers. Each box was provisioned with a cotton ball soaked in 10% honey solution. For 35 mate pairs that produced daughters, ovipositor length and bristle number were profiled for every mother and at least one and up to four of her daughters (mean n = 3.4). To interrogate bristle number independently of ovipositor length, we extracted residual bristle number from a linear regression of bristle number against ovipositor length; because the interaction between ovipositor length and generation (mother or daughters) was not significant, we included both generations in a single regression model. Narrow-sense heritability (*h^2^*) of residual ovipositor bristle number and ovipositor length – the proportion of phenotypic variation due to additive genetic effects – were each estimated by regressing the phenotype of each mother against the average phenotype of her daughters. Following convention when a trait can only be measured in parents of a single sex [7], *h^2^* was defined as twice the slope of the parent-offspring regression. A one-tailed *p* value was used to test the hypothesis that *h^2^* > 0.

### Read mapping and pool-GWA

Reads were trimmed using Trimmomatic v0.32 with the parameters “TRAILING:3 HEADCROP:2 SLIDINGWINDOW:6:15 MINLEN:50” and mapped to an *S. flava* reference genome assembly (GenBank accession no. GCA_003952975.1) using bwa v0.7.12 [8–10] with the following parameters: for bwa aln, “-o 3 –d 15 -l 100”; for bwa sampe, “- a 1000”. PCR and optical duplicate reads were removed using Picard Tools v1.107 (http://picard.sourceforge.net). Unpaired and low-quality reads were removed using the View command in Samtools v1.3.1 [11] with parameters “-q 20 –f 0x0002 –F 0x0004 –F 0x0008”. Low quality bases were removed using the Samtools mpileup command with parameters “-B –Q 17”. Repeat regions and 5 bp windows flanking indels (minimum count > 4) were filtered using Popoolation2 [12].

Statistical significance of allele frequency differences per site was estimated using the Cochran-Mantel-Haenszel test using Popoolation2 and custom scripts for sites with a minimum minor allele count of 10, coverage depth of 100, and minor allele frequency of 8% across all pools combined, and a minimum coverage depth of 15 and maximum coverage depth 200 in each pool.

We observed and adjusted for a minor observed inflation of *p* values. Systematic inflation of GWAS test statistics (termed genomic inflation) -- which is typically assumed to arise due to unmodeled relatedness among individuals, biased test implementation, or errors in genotyping -- can result in overly confident p-values. Genomic inflation is also expected to arise under polygenic control of a trait, even in the absence of population structure and other technical artefacts [13]. We identified and conservatively sought to correct for a slight inflation of p-values in our pool-GWA analysis [14]. However, because the distribution of p-values was non-uniform with an excess of both higher and lower values, typical corrections based on the observed vs. median test statistic gave unsuitable inflation factors. Following Thoen et al., we therefore regressed observed against expected –log_10_(*P*) values with the intercept constrained to 0, and divided each –log_10_(*P*) value by the slope of the regression line [15]. We excluded from our regression model the 1% most significant SNPs from our pool-GWAS (which are likely to be enriched for true associations) and SNPs that failed the stringent filtering described in our GO enrichment analysis (which may have overly conservative or liberal p-values due to biases or errors in genome assembly, sequencing and read mapping, and SNP genotyping, given the nature of our filters). Our approach for p-value adjustment was conservative, yielding a median p*-*value of 0.597.

### Linkage disequilibrium

#### Pooled genome sequencing

A total of 45 *S. flava* larvae were collected from *Turritis* (formerly *Arabis*) *glabra* from a large field (∼50,000 m^2^) in Belmont, MA, USA, which contained thousands of individual *T. glabra* plants with heavy *S. flava* mining damage, between June 22 and July 2, 2013. Each larva was collected from a separate plant individual to minimize relatedness. Samples were preserved in 95% ethanol at -80C. DNA was extracted from the pool of larvae using a DNeasy Blood and Tissue Kit (Qiagen). 100 bp paired-end sequencing was conducted on half of a lane on an Illumina HiSeq 2500 at the University of Arizona in January 2014.

#### Read mapping

Reads were trimmed of adapters, and trimmed and filtered for quality with Trimmomatic v. 0.32 [8] using the following settings: ILLUMINACLIP:2:30:10, TRAILING:3, HEADCROP:2, SLIDINGWINDOW:6:15, and MINLEN:50. Retained reads were then mapped to the *S. flava* reference genome first with BWA v.0.6.1 [9] using the MEM algorithm, and then using Stampy v.1.0.23 [16] using the -bamkeepgood reads option. From an initial Stampy run using a subset of the data, the substitution rate was obtained with Stampy and the average insert size was obtained with Picard v.1.107 CollectInsertSizeMetrics, and these estimates were used as parameters when mapping the full read set. Resulting SAM files were converted to BAM files using SAMtools v.0.1.18 [11]. BAM files were cleaned and sorted using Picard CleanSam and SortSam, and duplicate reads were marked and removed using Picard MarkDuplicates. Realignment around indels was performed using GATK v.2.8-1 [17] RealignerTargetCreator and IndelRealigner. SAMTools was then used to remove unmapped reads, keep only properly mapped read pairs, and filter for a mapping quality of 20. BEDTools v.2.17.0 [18] intersect was used to filter out repetitive regions, identified using the Drosophila repeat library in RepeatMasker v4.0.5. Reads were mapped to a mean coverage depth of 31.4x across the *S. flava* genome.

#### Linkage Disequilibrium estimates

LD was estimated from the 15 largest autosomal scaffolds. SNPs were called using GATK v.2.8-1 [17] UnifiedGenotyper with heterozygosity of 0.014, ploidy level of 90, two maximum alternative alleles, and a maximum coverage of 200. Preliminary SNPs were then hard filtered using GATK v.2.8-1 VariantFiltration and SelectVariants. LDx [19] was used to obtain a maximum likelihood estimate of linkage disequilibrium (r^2^) with the following parameters: a minimum read depth of 10, a maximum read depth of 150, an insert size of 417, a minimum quality score of 20, a minimum minor allele frequency of 0.1, and a minimum read intersection depth of 11.

### Gene ontology enrichment analysis on candidate SNPs

To test whether genes intersecting the top GWAS associations were enriched for particular predicted functions, we assigned functional annotations using orthology relationships among protein-coding genes in *S. flava* and other *Drosophila* genomes. Orthology was inferred by similarity clustering using orthoMCL v2.0.9 [20], with default parameters and an inflation value of 1.5, among proteomes for *S. flava,* all *Drosophila* species from the Drosophila 12 Genomes Consortium [21] (retrieved from FlyBase release 2013_06) except *D. willistoni*, and a draft genome assembly of *S. pallida* (unpublished data). Each *S. flava* gene was then annotated with the gene ontology (GO) terms assigned to its predicted ortholog(s) in *D. melanogaster* in FlyBase (release 2020_02). Parental GO terms that were implied but not directly listed, which were necessary for downstream analyses, were retrieved using GO.db v3.7.0 [22, 23].

We used GOWINDA [24] to test for enrichments of gene ontology (GO) terms among the genes that intersected SNPs with the strongest pool-GWAS associations (top 0.1% and 0.005% of p-values). We conservatively assumed that SNPs within the same gene were in LD and thus were not independent associations (--mode gene), and significance was determined from one million permutations. All gene models were extended by 200 bp (--gene-definition updownstream200) to account for the fact that genotyped SNPs may tag non-genotyped causal variants that are proximal, but the rapid decay of LD in *S. flava* (Fig. 4b) makes this unlikely over long physical distances. To capture both protein-coding and regulatory effects, SNPs were assigned to a given gene if they fell within its exons, introns, or the adjacent upstream or downstream intergenic region. Intergenic regions were extended from a focal gene’s UTR boundary until reaching the boundary of the adjacent gene’s UTR, up to a maximum of 2 kb. Intergenic regions were included because most cis-regulatory elements in *D. melanogaster* are located within or adjacent to the genes they regulate, but are rarely separated by an intervening gene [25]. To avoid diluting statistical power by the inclusion of redundant, nested GO terms or terms with few member genes, terms were only considered if they were assigned to at least 20 genes in *S. flava,* and we focused on only a single level of the GO hierarchy (where level refers to the number of edges from the focal GO term to the root of the acyclic graph of GO term relationships). Level two was used for Molecular Function, which maximized the number of terms considered, and level four was used for Biological Process because it contained a similar number of terms.

Prior to GO enrichment analyses, we performed a more stringent SNP filtering step, excluding tri-allelic sites (having a frequency > 0.08 for the third most common allele) and SNPs located within 300 bp of a scaffold edge.

### Replication of a candidate SNP upstream of *Gai*

DNA was extracted from whole adult flies with the Qiagen DNeasy blood and tissue kit, following the provided protocol. PCR primers for the *Gαi* region were designed in Geneious, using default settings and a target region size of 500bp. The primer sequences are as follows: gai4F: CATTTCTGTCCATGGCGTCG; gai9R: GCCGTTAGACAAAGCGCATT. PCR methods were the same as those in Gloss et al. (2013). PCR clean up and sequencing were performed at the UC Berkeley DNA Sequencing Facility.

For trimming, alignment, and base calling, we used Geneious v.10.0.5 (Biomatters Ltd.) Using the Trimming Tool in Geneious, regions with more than a 5% chance of an error per base were trimmed. Sequences were aligned with default “Geneious Alignment” setting (Cost matrix: 65% similarity, Gap open penalty: 12, Gap extension penalty: 3, Refinement Iterations: 2, Alignment type: Global alignment with free end gaps). The final alignment was 331 base pairs long. A significant fraction of sequences had convoluted regions upstream of the SNP, likely due to heterozygous indels. Manual base-calling was therefore performed on all sequences. Double peaks were called as heterozygotes. Sequences with convoluted regions were analyzed with Indelligent v.1.2, which identified indels [26]. In these cases, most of the reverse strand sequence was rendered unreadable, so all sequences (convoluted or not) were based on only forward strand nucleotide base calls. Variant sites, including the candidate SNP, were identified in Geneious, using a minimum variant frequency cutoff of 0.05.

We performed a test for Hardy-Weinberg equilibrium (HWE) for the focal SNP among all individuals using the *hw.test* function from the pegas package. The SNP was found to be in HWE (χ2 = 3.15, P > 0.05).

## Supplemental Figures

**Figure S1.**
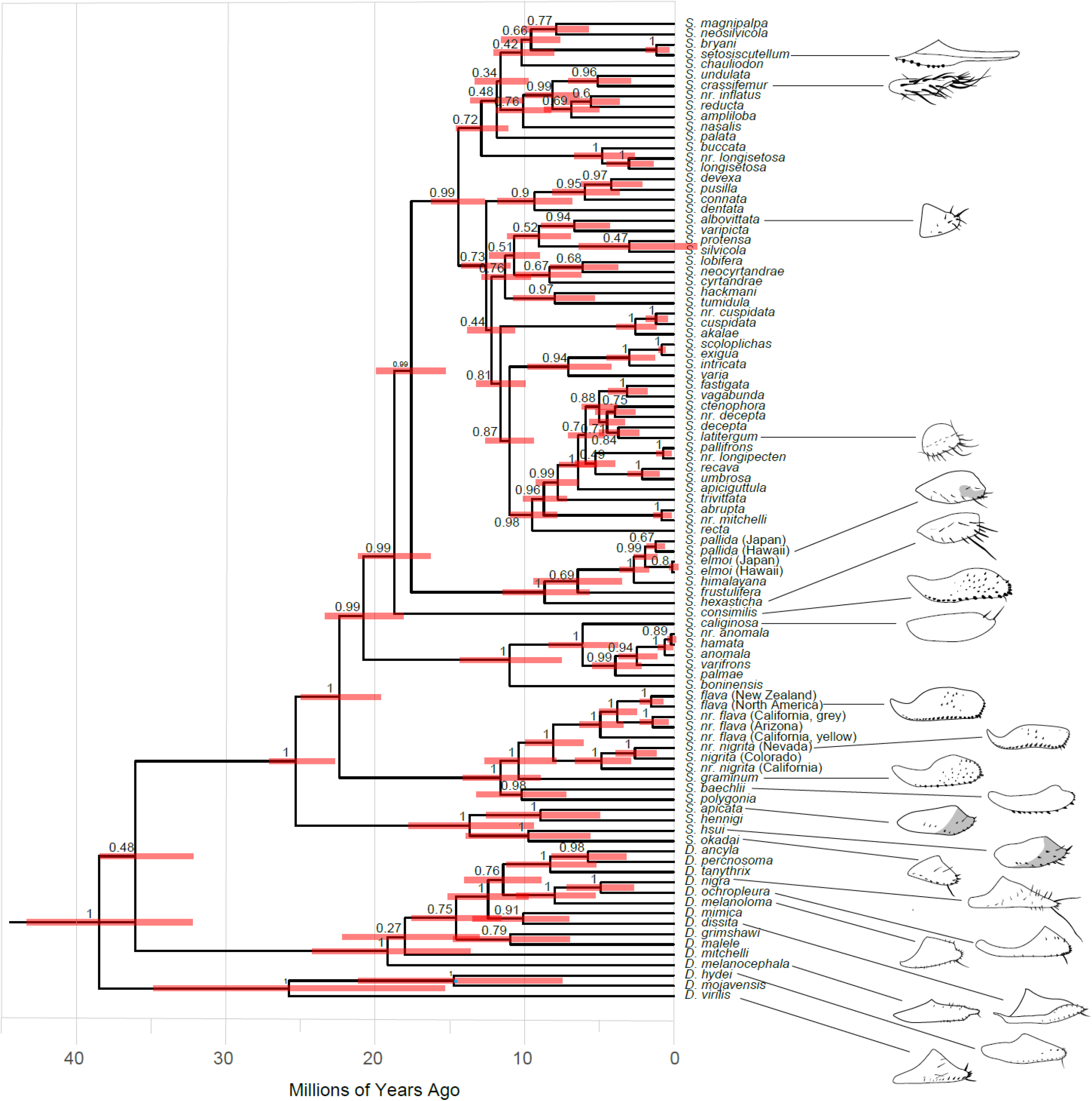
Bayesian tree. **Time calibrated phylogeny of subgenus *Drosophila* and *Scaptomyza* inferred from Bayesian analysis**, using 11 genes (16S, 28S, *Adh*, *Cad-r* (rudimentary), *COI*, *COII*, *gstd1*, *gpdh*, *marf*, *ND2*, *n(l)tid)*, *Orco*) and fossil and biogeographic time calibrations. Nodes indicate posterior probabilities, red bars represent 95% highest posterior density interval around the mean node age. Illustrations of female ovipositors, indicating bristle number, were drawn from sources indicated in Table S1.

**Figure S2.**
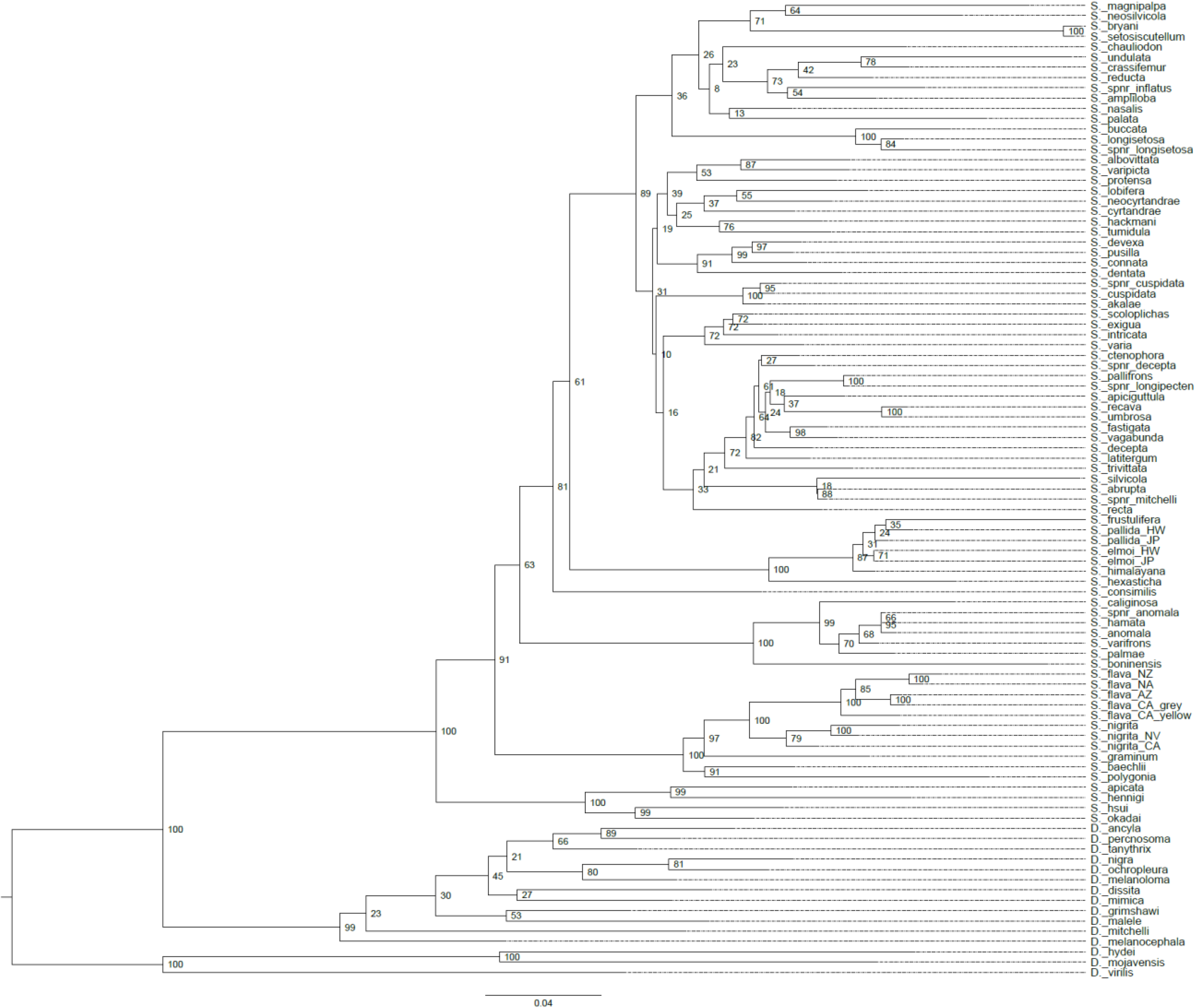
ML tree. **Phylogeny of subgenus *Drosophila* and *Scaptomyza* inferred from maximum likelihood (ML),** using 11 genes (16S, 28S, *Adh*, *Cad-r* (rudimentary), *COI*, *COII*, *gstd1*, *gpdh*, *marf*, *ND2*, *n(l)tid)*, *Orco*) and fossil and biogeographic time calibrations. Nodes represent bootstrap value (≥50%) from ML analysis.

**Figure S3.**
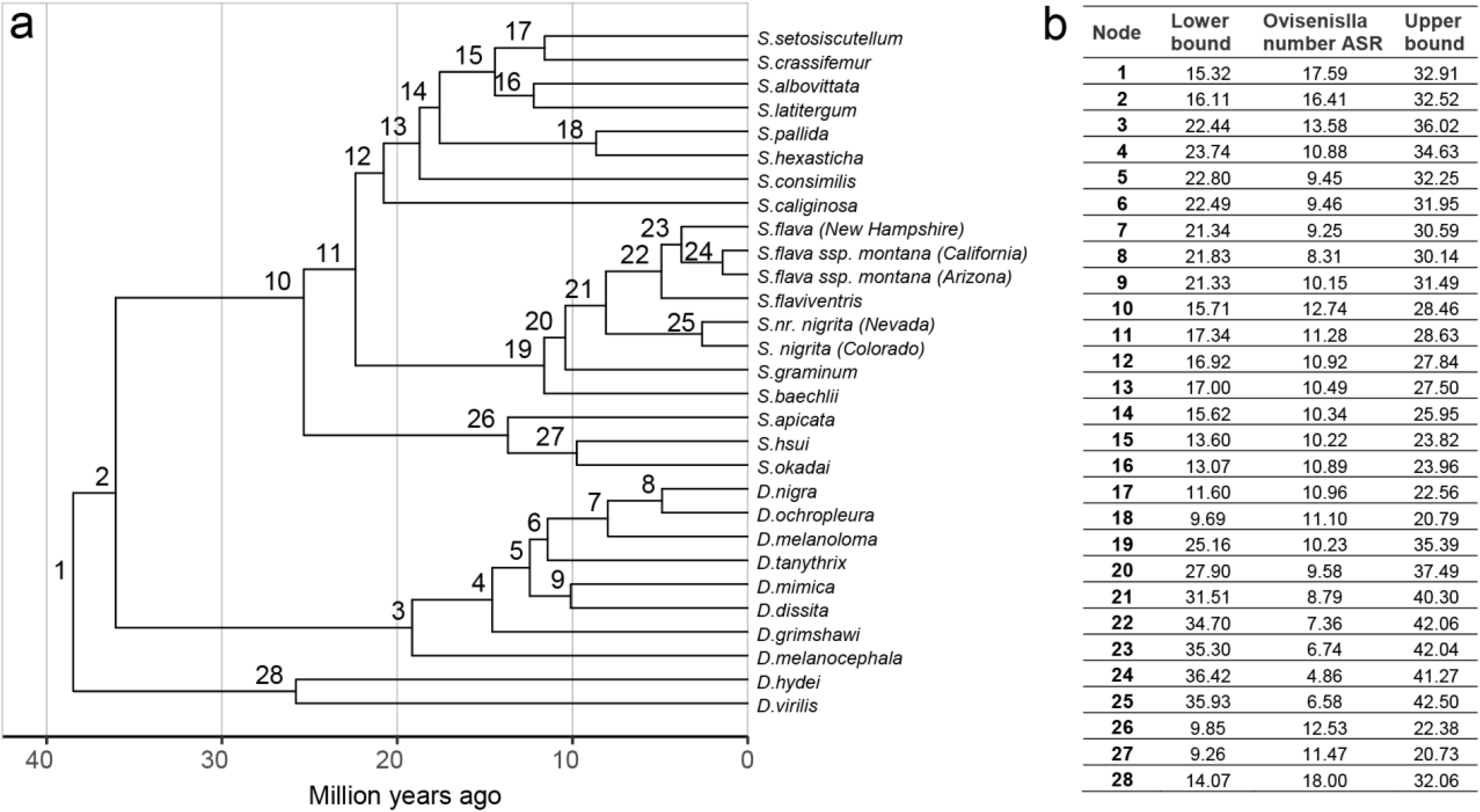
CI for ASR on ovipositor bristle number. Confidence intervals for ancestral state estimations of ovipositor bristle number. **Values in** the table correspond to respective nodes given in the phylogeny.

**Figure S4.**
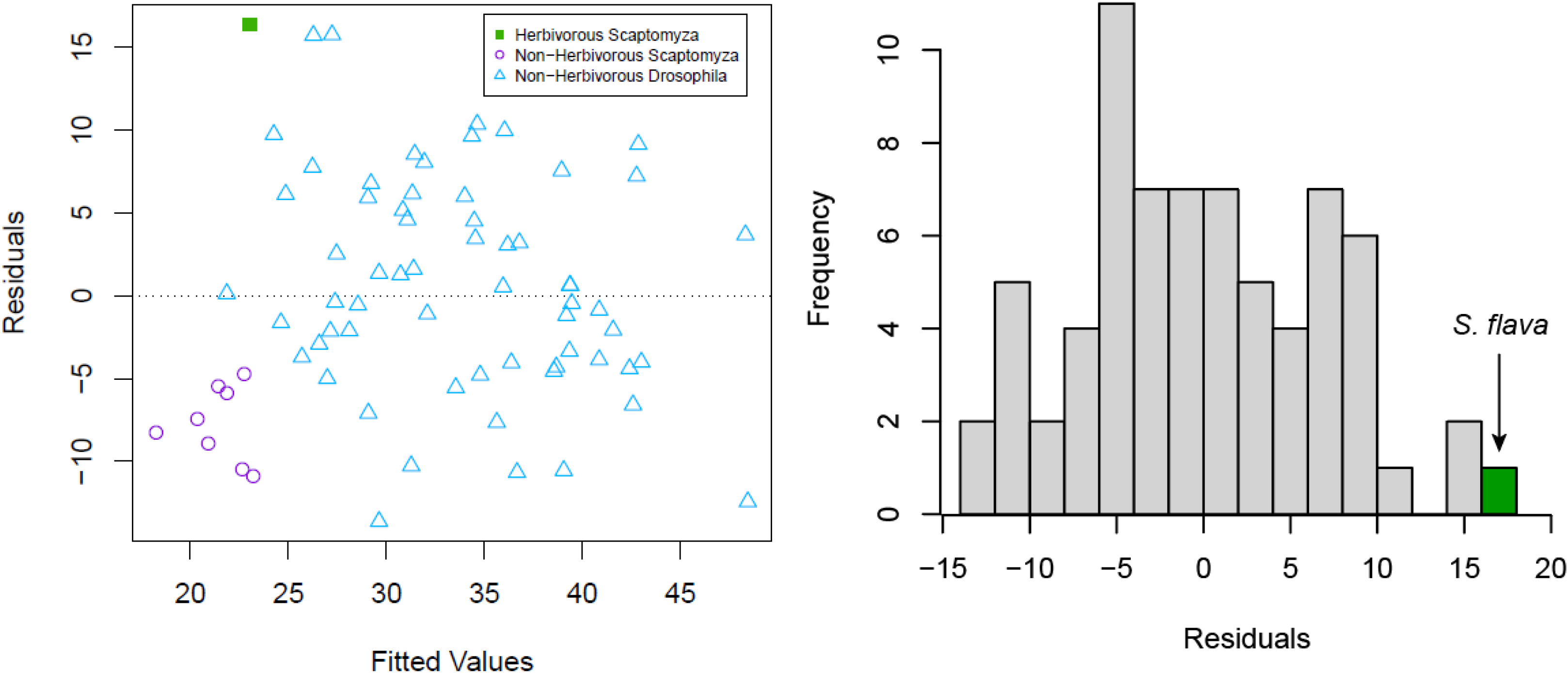
Residual Analysis. Residual analysis on linear regression model on ovipositor bristle number. Ovipositor bristle number, ovipositor length, and larval feeding ecology were obtained from Craddock et al. 2018, with additional data from this study (Suppl. Dataset S3). (a) Scatter plot of residuals on the y-axis and fitted values (estimated responses) on the x axis. (b) Histogram of residuals from (a).

## Supplemental Tables

**Table S1.**
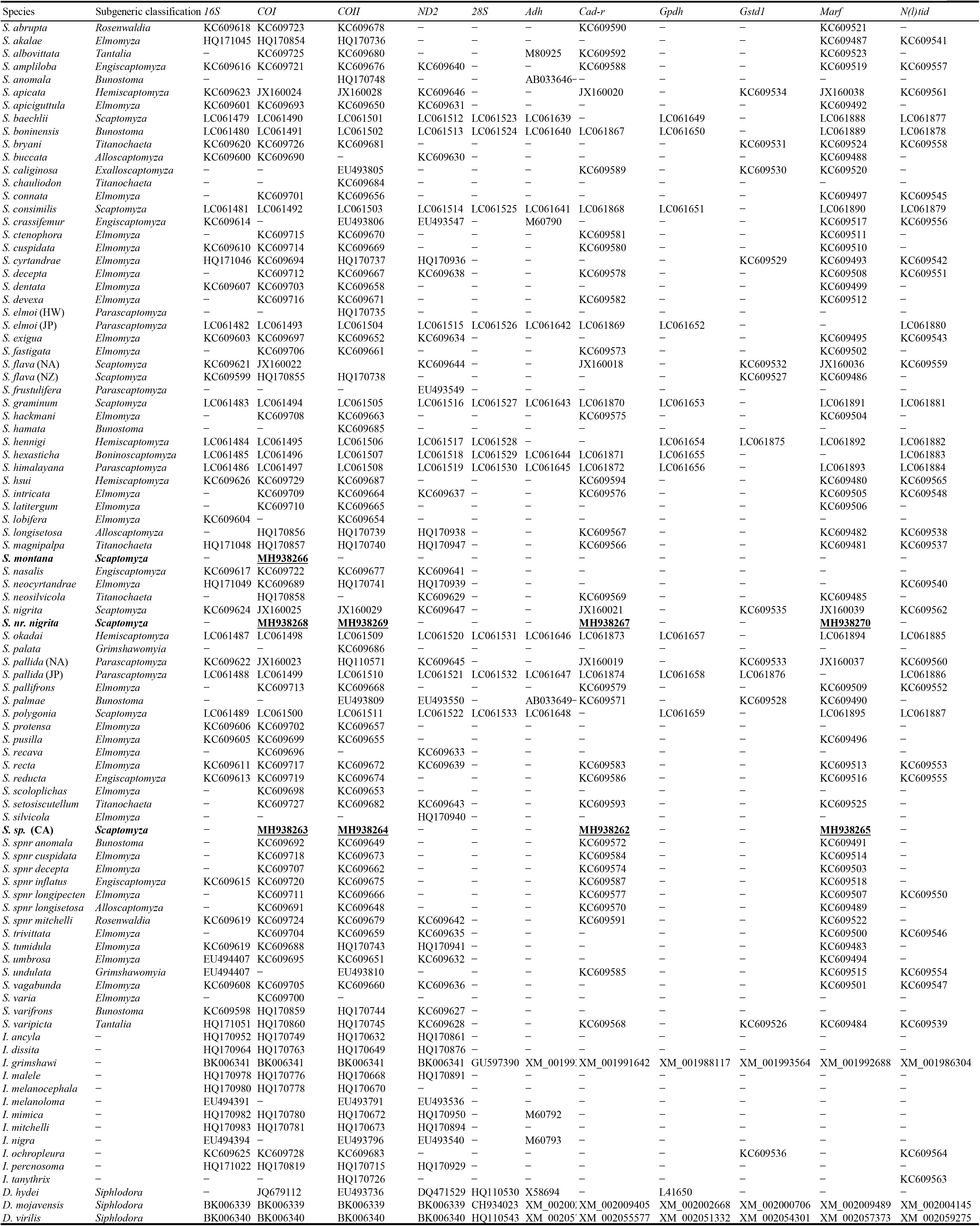
Species included in this study and GenBank accession numbers for sequences included. Sequences obtained in this study are in underlined bold font.

**Table S2.** Ovipositor bristle number and larval feeding ecology data for 95 species included in phylogenetic analyses to estimate the ancestral character states and perform phylogenetic generalized least squares. * Data from Lapoint et al. 2013.

| Species | Larval Diet* | Source for Bristle Counts | Ovipositor Bristles |
| --- | --- | --- | --- |
| *D. ancyla* | saprophagy |  |  |
| *D. dissita* | saprophagy | Throckmorton 1966 | 18 |
| *D. grimshawi* | saprophagy | Craddock et al. 2018 | 36,42, 43 |
| *D. hydei* | saprophagy | Baechli et al. 2004 | 23 |
| *D. malele* | unknown |  |  |
| *D. melanocephala* | saprophagy | Craddock et al. 2018 | 39 |
|  |  | Hardy 1967 | 31 |
| *D. melanoloma* | fungus feeding | Throckmorton 1966 | 23 |
| *D. mimica* | saprophagy | Craddock et al. 2018 | 30, 30 |
| *D. mitchelli* | unknown |  |  |
| *D. mojavenensis* | saprophagy |  |  |
| *D. nigra* | unknown | Hardy et al. 2001 | 28 |
|  |  | Throckmorton 1966 | 27 |
|  |  | Craddock et al. 2018 | 33, 33 |
| *D. ochropleura* | fungus feeding | Hardy et al. 2001 | 21 |
|  |  | Craddock et al. 2018 | 19 |
|  |  | Craddock et al. 2018 | 23 |
|  |  | Craddock et al. 2018 | 24 |
| *D. percnosoma* | saprophagy |  |  |
| *D. tanythrix* | saprophagy | Craddock et al. 2018 | 28 |
| *D. virilis* | saprophagy | Sturtevant 1921 | 20 |
| *S. abrupta* | unknown |  |  |
| *S. akalae* | unknown |  |  |
| *S. albovittata* | unknown | Craddock & Kambysellis 1997 | 12 |
|  |  | Craddock et al. 2018 | 10 |
| *S. ampliloba* | unknown |  |  |
| *S. anomala* | unknown |  |  |
| *S. nr. anomala* | unknown |  |  |
| *S. apicata* | saprophagy | Mount Hood National Forest,<br>Camp Creek | 11 |
|  |  | Mount Hood National Forest,<br>Trout Creek | 13, 13 |
|  |  | Lab colony reared from wild<br>caught individuals from<br>Strawberry Creek, Berkeley,<br>California | 12, 14, 15 |
| *S. apiciguttula* | saprophagy |  |  |
| *S. baechlii* | unknown | Sidorenko 1993 | 15 |
| *S. boninensis* | unknown | Okada 1973 |  |
| *S. bryani* | spider predation |  |  |
| *S. buccata* | unknown |  |  |
| *S. caliginosa* | flower specialist | Craddock et al. 2018 | 16 |
| *S. chauliodon* | spider predation |  |  |
| *S. connata* | unknown |  |  |
| *S. consimilis* | saprophagy | Baechli et al. 2004 | 45 |
| *S. crassifemur* | unknown | Throckmorton 1966 | 20 |
|  |  | Craddock et al. 2018 | 18 |
|  |  | Grimaldi | 20 |
|  |  | Hardy 1965 | 20 |
| *S. ctenophora* | unknown |  |  |
| *S. cuspidata* | saprophagy |  |  |
| *S. nr. cuspidata* | saprophagy |  |  |
| *S. cyrtandrae* | saprophagy |  |  |
| *S. decepta* | unknown |  |  |
| *S. nr. decepta* | unknown |  |  |
| *S. dentata* | unknown |  |  |
| *S. devexa* | unknown |  |  |
| *S. elmoi* (Hawaii) | saprophagy |  |  |
| *S. elmoi* (Japan) | saprophagy |  |  |
| *S. exigua* | saprophagy |  |  |
| *S. fastigata* | unknown |  |  |
| *S. flava* (North America) | leaf-mining | Lab colony, reared from wild caught individuals from New Hampshire | 34, 35, 38, 38, 40, 40, 40, 40, 41, 41, 41, 44 |
| *S. flava* (New Zealand) | leaf-mining |  |  |
| *S. flava ssp. montana* (California) | leaf-mining | Strawberry Creek, Berkeley, California | 37, 39 |
|  |  | Lab colony, reared from wild caught individuals from Strawberry Creek, Berkeley, CA | 27, 36, 37, 37, 38, 40, 40, 42 |
| *S. flava ssp. montana* (Arizona) | leaf-mining | Lab colony, reared from wild caught individuals from Arizona | 37, 37, 37, 39, 40, 41, 42, 42, 44, 46 |
| *S. flaviventris* | leaf-mining | Santa Cruz, California | 39, 43 |
| *S. frustulifera* | unknown |  |  |
| *S. graminum* | leaf-mining | Baechli et al. 2004 | 39 |
| *S. hackmani* | saprophagy |  |  |
| *S. hamata* | unknown |  |  |
| *S. hennigi* | unknown |  |  |
| *S. hexasticha* | unknown | Okada 1973 | 14 |
| *S. himalayana* | unknown |  |  |
| *S. hsui* | saprophagy | Thomas Creek, Nevada | 11, 11, 12, 12, 12, 12, 13 |
|  |  | Mount Hood National Forest, Still Creek, Oregon | 12 |
|  |  | Rocky Mountain Biological Lab, near Gothic, Colorado | 12, 15 |
| *S. nr. inflatus* | unknown |  |  |
| *S. intricata* | saprophagy |  |  |
| *S. latitergum* | saprophagy | Throckmorton 1966 | 25 |
| *S. lobifera* | unknown |  |  |
| *S. nr. longipecten* | saprophagy |  |  |
| *S. longisetosa* | unknown |  |  |
| *S. nr. longisetosa* | unknown |  |  |
| *S. magnipalpa* | spider predation |  |  |
| *S. nr. mitchelli* | unknown |  |  |
| *S. nasalis* | unknown |  |  |
| *S. neocyrtandrae* | saprophagy |  |  |
| *S. neosilvicola* | spider predation |  |  |
| *S. nigrita* (Colorado) | leaf-mining | Rocky Mountain Biological Lab,<br>near Gothic, Colorado | 37, 38, 39, 41, 41, 42,<br>42 |
| *S. nr. nigrita*<br>(California) | leaf-mining |  |  |
| *S. nr. nigrita*<br>(Nevada) | leaf-mining | Lab colony, reared from wild<br>caught individuals from Lake<br>Tahoe, Nevada | 38, 38, 44 |
| *S. okadai* | unknown | Hackman 1959 | 15 |
| *S. palata* | unknown |  |  |
| *S. pallida* (Japan) | saprophagy |  |  |
| *S. pallida* (North<br>American) | saprophagy | Mount Hood National Forest, Still<br>Creek, Oregon | 8, 8, 9, 10, 11, 12, 12 |
|  |  | Mount Hood National Forest,<br>Camp Creek, Oregon | 13 |
|  |  | Baechli et al. 2004 | 17 |
| *S. pallifrons* | unknown |  |  |
| *S. palmae* | flower specialist |  |  |
| *S. polygonia* | saprophagy |  |  |
| *S. protensa* | unknown |  |  |
| *S. pusilla* | unknown |  |  |
| *S. recava* | unknown |  |  |
| *S. recta* | unknown |  |  |
| *S. reducta* | unknown |  |  |
| *S. scoloplichas* | saprophagy |  |  |
| *S. setosiscutellum* | spider predation | Hardy 1965 | 8 |
| *S. silvicola* | unknown |  |  |
| *S. trivittata* | unknown |  |  |
| *S. tumidula* | saprophagy |  |  |
| *S. umbrosa* | unknown |  |  |
| *S. undulata* | unknown |  |  |
| *S. vagabunda* | unknown |  |  |
| *S. varia* | saprophagy |  |  |
| *S. varipicta* | saprophagy |  |  |
| *S. varifrons* | unknown |  |  |

**Table S3.**
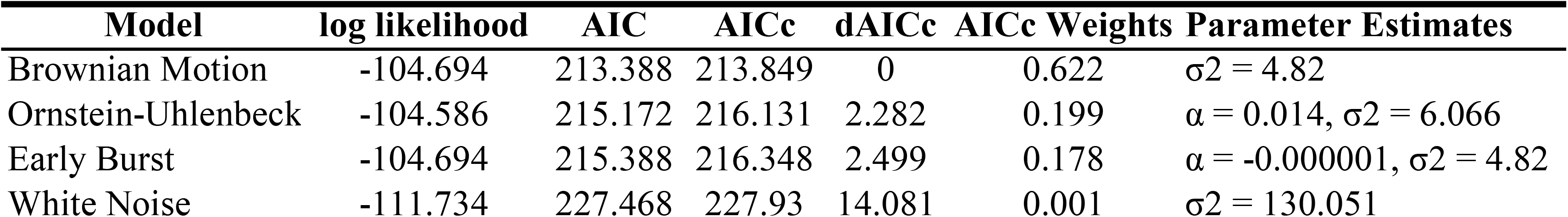
Comparison of models of evolution for ovipositor bristle number.

**Table S4.**
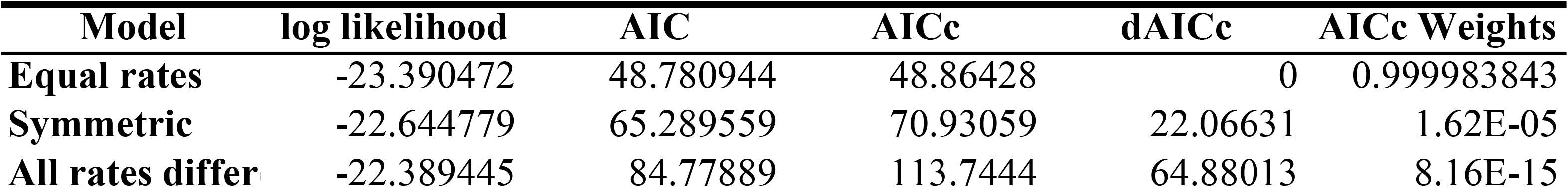
Comparison of models of evolution for larval diet.

**Table S5.**
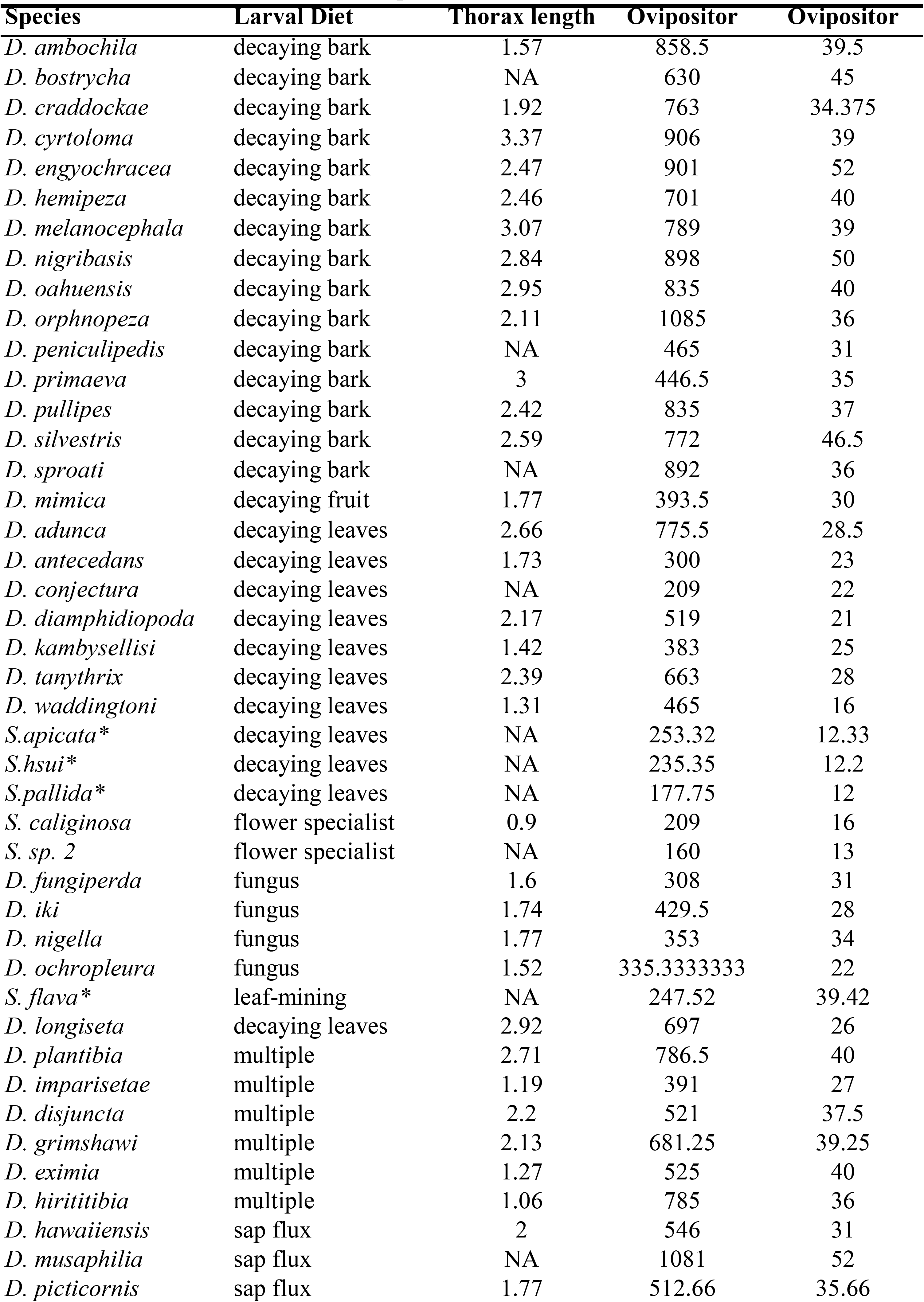
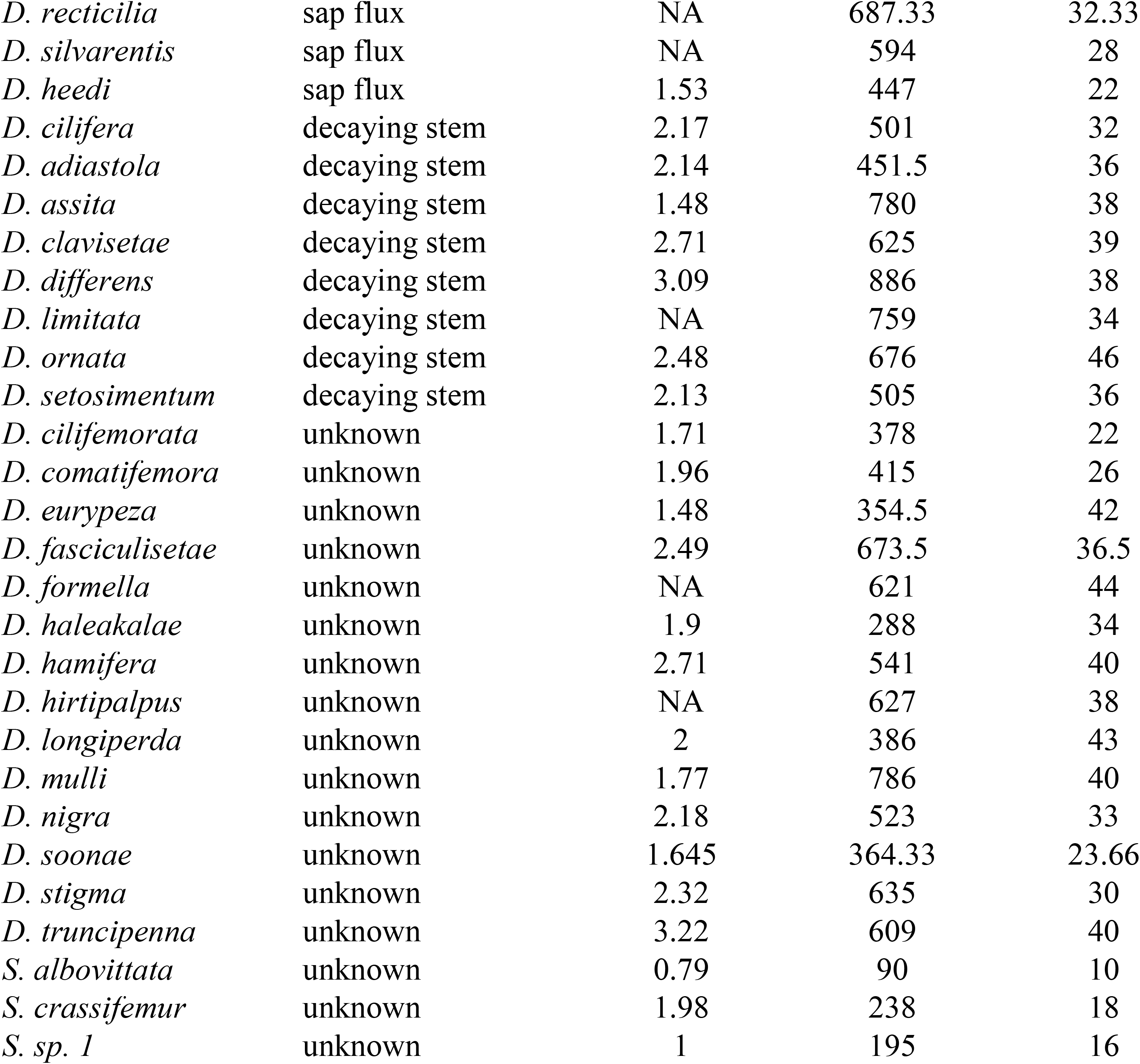
Ovipositor length, ovipositor bristle number, and larval feeding ecology data for 67 species from Craddock et al. 2018 and 4 species from this study (denoted with *), used to evaluate the relationship between ovipositor length and bristle number. Ovipositor bristle counts were averaged across individuals and ovipositor valves when measures for both were present.

| Species | Larval Diet | Thorax length | Ovipositor | Ovipositor |
| --- | --- | --- | --- | --- |
| *D. ambochila* | decaying bark | 1.57 | 858.5 | 39.5 |
| *D. bostrycha* | decaying bark | NA | 630 | 45 |
| *D. craddockae* | decaying bark | 1.92 | 763 | 34.375 |
| *D. cyrtoloma* | decaying bark | 3.37 | 906 | 39 |
| *D. engyochracea* | decaying bark | 2.47 | 901 | 52 |
| *D. hemipeza* | decaying bark | 2.46 | 701 | 40 |
| *D. melanocephala* | decaying bark | 3.07 | 789 | 39 |
| *D. nigribasis* | decaying bark | 2.84 | 898 | 50 |
| *D. oahuensis* | decaying bark | 2.95 | 835 | 40 |
| *D. orphnopeza* | decaying bark | 2.11 | 1085 | 36 |
| *D. peniculipedis* | decaying bark | NA | 465 | 31 |
| *D. primaeva* | decaying bark | 3 | 446.5 | 35 |
| *D. pullipes* | decaying bark | 2.42 | 835 | 37 |
| *D. silvestris* | decaying bark | 2.59 | 772 | 46.5 |
| *D. sproati* | decaying bark | NA | 892 | 36 |
| *D. mimica* | decaying fruit | 1.77 | 393.5 | 30 |
| *D. adunca* | decaying leaves | 2.66 | 775.5 | 28.5 |
| *D. antecedans* | decaying leaves | 1.73 | 300 | 23 |
| *D. conjectura* | decaying leaves | NA | 209 | 22 |
| *D. diamphidiopoda* | decaying leaves | 2.17 | 519 | 21 |
| *D. kambysellisi* | decaying leaves | 1.42 | 383 | 25 |
| *D. tanythrix* | decaying leaves | 2.39 | 663 | 28 |
| *D. waddingtoni* | decaying leaves | 1.31 | 465 | 16 |
| *S. apicata* * | decaying leaves | NA | 253.32 | 12.33 |
| *S. hsui* * | decaying leaves | NA | 235.35 | 12.2 |
| *S. pallida* * | decaying leaves | NA | 177.75 | 12 |
| *S. caliginosa* | flower specialist | 0.9 | 209 | 16 |
| *S. sp. 2* | flower specialist | NA | 160 | 13 |
| *D. fungiperda* | fungus | 1.6 | 308 | 31 |
| *D. iki* | fungus | 1.74 | 429.5 | 28 |
| *D. nigella* | fungus | 1.77 | 353 | 34 |
| *D. ochropleura* | fungus | 1.52 | 335.3333333 | 22 |
| *S. flava* * | leaf-mining | NA | 247.52 | 39.42 |
| *D. longiseta* | decaying leaves | 2.92 | 697 | 26 |
| *D. plantibia* | multiple | 2.71 | 786.5 | 40 |
| *D. imparisetae* | multiple | 1.19 | 391 | 27 |
| *D. disjuncta* | multiple | 2.2 | 521 | 37.5 |
| *D. grimshawi* | multiple | 2.13 | 681.25 | 39.25 |
| *D. eximia* | multiple | 1.27 | 525 | 40 |
| *D. hirititibia* | multiple | 1.06 | 785 | 36 |
| *D. hawaiiensis* | sap flux | 2 | 546 | 31 |
| *D. musaphilia* | sap flux | NA | 1081 | 52 |
| *D. picticornis* | sap flux | 1.77 | 512.66 | 35.66 |
| *D. recticilia* | sap flux | NA | 687.33 | 32.33 |
| *D. silvarentis* | sap flux | NA | 594 | 28 |
| *D. heedi* | sap flux | 1.53 | 447 | 22 |
| *D. cilifera* | decaying stem | 2.17 | 501 | 32 |
| *D. adiaistola* | decaying stem | 2.14 | 451.5 | 36 |
| *D. assita* | decaying stem | 1.48 | 780 | 38 |
| *D. clavisetae* | decaying stem | 2.71 | 625 | 39 |
| *D. differens* | decaying stem | 3.09 | 886 | 38 |
| *D. limitata* | decaying stem | NA | 759 | 34 |
| *D. ornata* | decaying stem | 2.48 | 676 | 46 |
| *D. setosimentum* | decaying stem | 2.13 | 505 | 36 |
| *D. cilifemorata* | unknown | 1.71 | 378 | 22 |
| *D. comatifemora* | unknown | 1.96 | 415 | 26 |
| *D. eurypeza* | unknown | 1.48 | 354.5 | 42 |
| *D. fasciculisetae* | unknown | 2.49 | 673.5 | 36.5 |
| *D. formella* | unknown | NA | 621 | 44 |
| *D. haleakalae* | unknown | 1.9 | 288 | 34 |
| *D. hamifera* | unknown | 2.71 | 541 | 40 |
| *D. hirtipalpus* | unknown | NA | 627 | 38 |
| *D. longiperda* | unknown | 2 | 386 | 43 |
| *D. mulli* | unknown | 1.77 | 786 | 40 |
| *D. nigra* | unknown | 2.18 | 523 | 33 |
| *D. soonae* | unknown | 1.645 | 364.33 | 23.66 |
| *D. stigma* | unknown | 2.32 | 635 | 30 |
| *D. truncipenna* | unknown | 3.22 | 609 | 40 |
| *S. albovittata* | unknown | 0.79 | 90 | 10 |
| *S. crassifemur* | unknown | 1.98 | 238 | 18 |
| *S. sp. 1* | unknown | 1 | 195 | 16 |

**Table S6.** Full dataset used to evaluate whether SNP effect sizes identified from the pool-GWAS on ovipositor bristle number in *S. flava* could be replicated through individual sequencing. SNP214 represents the candidate SNP identified from the pool-GWAS. Other SNPs and indels that were present at a minimum variant frequency of 0.05 within the sequenced region are also presented. Ovipositor length is given in um. Raw sequence data is available on Genbank (accession no. MH884655-MH884734).

| ID | Population_HostPlant | Population | PegPool | PegNumConsensus | OviLengthConsensus | SNP214 | SNP214. | SNP49 | Indel213 | SNP215 | SNP235 | SNP236 |
| --- | --- | --- | --- | --- | --- | --- | --- | --- | --- | --- | --- | --- |
| P4RAC11 | NH1_TurrBarb | NH1 | low | 35 | 0.2517 | AA | 0 | AA | TT | TT | TT | CC |
| P4RAC8 | NH1_TurrBarb | NH1 | high | 42 | 0.25407 | CA | 1 | AA | T- | GG | NA | NA |
| P4RBC1 | NH1_TurrBarb | NH1 | high | 44 | 0.24112 | CC | 2 | TT | -- | GG | CC | TT |
| P4RBC2 | NH1_TurrBarb | NH1 | low | 34 | 0.259545 | AA | 0 | AA | TT | TT | TT | CC |
| P4RBC4 | NH1_TurrBarb | NH1 | high | 45 | 0.2343275 | CC | 2 | TT | -- | GG | CC | TT |
| P4RBC5 | NH1_TurrBarb | NH1 | low | 35 | 0.23715 | AA | 0 | AA | TT | TT | TT | CC |
| P4RBC6 | NH1_TurrBarb | NH1 | high | 46 | 0.2461875 | CA | 1 | AA | T- | GG | NA | NA |
| P4RBC7 | NH1_TurrBarb | NH1 | high | 41 | 0.23348 | CC | 2 | AA | -- | TT | CC | TT |
| P4RCC10 | NH1_TurrBarb | NH1 | high | 44 | 0.2679325 | CC | 2 | TT | -- | GG | CC | TT |
| P4RCC11 | NH1_TurrBarb | NH1 | high | 42 | 0.2499625 | CA | 1 | AA | T- | TT | TT | CC |
| P4RCC12 | NH1_TurrBarb | NH1 | high | 44 | 0.2506275 | CC | 2 | TT | -- | GG | CC | TT |
| P4RCC3 | NH1_TurrBarb | NH1 | high | 44 | 0.26938 | AA | 0 | AA | TT | TT | TT | CC |
| P4RCC9 | NH1_TurrBarb | NH1 | high | 42 | 0.23166 | CA | 1 | AA | TT | GT | CT | CT |
| P4RDC1 | NH1_TurrBarb | NH1 | low | 36 | 0.249805 | CC | 2 | AA | TT | GG | CC | TT |
| P4REC3 | NH1_TurrBarb | NH1 | low | 34 | 0.2359825 | AA | 0 | AA | TT | TT | TT | CC |
| P4REC6 | NH1_TurrBarb | NH1 | high | 43 | 0.26402 | CC | 2 | TT | -- | GG | CC | TT |
| P4REC7 | NH1_TurrBarb | NH1 | low | 34 | 0.2461375 | AA | 0 | AA | TT | TT | TT | CC |
| P4REC8 | NH1_TurrBarb | NH1 | low | 34 | 0.2495625 | CC | 2 | AA | -- | GG | CC | TT |
| P4RFC1 | NH1_TurrBarb | NH1 | low | 34 | 0.228205 | CA | 1 | AA | T- | TT | TT | CC |
| P4RFC11 | NH1_TurrBarb | NH1 | low | 37 | 0.26311 | CC | 2 | TT | -- | GG | CC | TT |
| P4RFC3 | NH1_TurrBarb | NH1 | low | 37 | 0.2603725 | CA | 1 | AA | T- | TT | TT | CC |
| P4RGC2 | NH1_TurrBarb | NH1 | high | 44 | 0.2702375 | CC | 2 | TT | -- | GG | CC | TT |
| P4RGC4 | NH1_TurrBarb | NH1 | high | 42 | 0.2425425 | AA | 0 | AA | TT | TT | TT | CC |
| P4RGC5 | NH1_TurrBarb | NH1 | high | 43 | 0.236325 | CC | 2 | TT | -- | GG | CC | TT |
| P4RGC8 | NH1_TurrBarb | NH1 | low | 34 | 0.24738 | AA | 0 | AA | TT | TT | TT | CC |
| P4RGC9 | NH1_TurrBarb | NH1 | low | 34 | 0.245095 | CA | 1 | AA | T- | TT | TT | CC |
| P4RHC1 | NH1_TurrBarb | NH1 | low | 36 | 0.267935 | CA | 1 | AA | T- | GG | NA | NA |
| P4RHC11 | NH1_TurrBarb | NH1 | high | 46 | 0.25933 | CC | 2 | TT | -- | GG | CC | TT |
| P4RHC3 | NH1_TurrBarb | NH1 | high | 44 | 0.2554425 | CA | 1 | AA | T- | TT | TT | CC |
| P4RHC4 | NH1_TurrBarb | NH1 | low | 33 | 0.25277 | AA | 0 | AA | TT | TT | TT | CC |
| P4RHC6 | NH1_TurrBarb | NH1 | high | 45 | 0.25376 | AA | 0 | AA | TT | TT | TT | CC |
| P5RAC4 | NH1_TurrBarb | NH1 | low | 37 | 0.2677875 | CC | 2 | TT | -- | GG | CC | TT |
| P5RAC6 | NH1_TurrBarb | NH1 | low | 35 | 0.243765 | AA | 0 | AA | TT | TT | TT | CC |
| P5RBC1 | NH1_TurrBarb | NH1 | low | 33 | 0.243675 | AA | 0 | AA | TT | TT | TT | CC |
| P5RBC10 | NH1_TurrBarb | NH1 | low | 37 | 0.2280675 | CC | 2 | AA | -- | GG | CC | TT |
| P5RBC2 | NH1_TurrBarb | NH1 | high | 46 | 0.2396925 | CC | 2 | TT | -- | GG | CC | TT |
| P5RBC3 | NH1_TurrBarb | NH1 | low | 33 | 0.250805 | CA | 1 | AA | T- | GG | NA | NA |
| P6RAC10 | NH2_Turr | NH2 | high | 45 | 0.2258725 | CA | 1 | AA | T- | TT | NA | NA |
| P6RAC11 | NH2_Turr | NH2 | low | 35 | 0.2362175 | CA | 1 | AA | T- | TT | NA | NA |
| P6RAC4 | NH2_Turr | NH2 | low | 38 | 0.259065 | CC | 2 | AA | TT | GG | CC | TT |
| P6RAC5 | NH2_Turr | NH2 | low | 39 | 0.2686325 | CA | 1 | AA | T- | GG | NA | NA |
| P6RBC11 | NH2_Turr | NH2 | low | 37 | 0.254155 | CC | 2 | TT | -- | GG | CC | TT |
| P6RBC2 | NH2_Turr | NH2 | high | 46 | 0.2667175 | CC | 2 | AA | -- | GG | CC | TT |
| P6RBC3 | NH2_Turr | NH2 | high | 43 | 0.2447975 | CC | 2 | AA | TT | GG | CC | TT |
| P6RBC4 | NH2_Turr | NH2 | high | 45 | 0.2592575 | CC | 2 | TT | -- | GG | CC | TT |
| P6RBC7 | NH2_Turr | NH2 | low | 33 | 0.2423825 | CA | 1 | AA | TT | GT | CT | CC |
| P6RCC1 | NH2_Turr | NH2 | low | 37 | 0.2520475 | CC | 2 | AA | -- | GG | CC | TT |
| P6RDC1 | NH2_Turr | NH2 | low | 37 | 0.2641875 | CA | 1 | AA | T- | TT | NA | NA |
| P6RDC11 | NH2_Turr | NH2 | high | 41 | 0.24728 | AA | 0 | AA | TT | TT | TT | CC |
| P6RDC4 | NH2_Turr | NH2 | high | 43 | 0.2496975 | CC | 2 | TT | -- | GG | CC | TT |
| P6RDC5 | NH2_Turr | NH2 | low | 36 | 0.2554225 | CA | 1 | AA | T- | TT | NA | NA |
| P6REC4 | NH2_Turr | NH2 | high | 44 | 0.254275 | CC | 2 | AA | -- | GG | CC | TT |
| P6REC5 | NH2_Turr | NH2 | high | 42 | 0.25232 | AA | 0 | AA | TT | TT | TT | CC |
| P6REC8 | NH2_Turr | NH2 | low | 37 | 0.264415 | AA | 0 | AA | TT | TT | TT | CC |
| P7RAC1 | NH2_Barb | NH2 | high | 43 | 0.26528 | CC | 2 | TT | -- | GG | CC | TT |
| P7RAC11 | NH2_Barb | NH2 | high | 45 | 0.2556975 | CA | 1 | AA | T- | GG | NA | NA |
| P7RAC5 | NH2_Barb | NH2 | high | 43 | 0.26603 | CC | 2 | AA | -- | GG | CC | TT |
| P7RAC6 | NH2_Barb | NH2 | low | 35 | 0.2552775 | AA | 0 | AA | TT | TT | TT | CC |
| P7RAC8 | NH2_Barb | NH2 | high | 43 | 0.2599025 | CC | 2 | AA | -- | GT | CC | TT |
| P7RBC1 | NH2_Barb | NH2 | low | 34 | 0.23766 | CA | 1 | AA | T- | TT | TT | CC |
| P7RBC10 | NH2_Barb | NH2 | high | 41 | 0.2358675 | CC | 2 | AA | -- | TT | CC | TT |
| P7RBC2 | NH2_Barb | NH2 | low | 36 | 0.259495 | CA | 1 | AA | T- | GG | NA | NA |
| P7RBC3 | NH2_Barb | NH2 | high | 41 | 0.24551 | CC | 2 | TT | -- | GG | CC | TT |
| P7RBC6 | NH2_Barb | NH2 | high | 44 | 0.264935 | CA | 1 | AA | TT | GT | TT | CC |
| P7RCC6 | NH2_Barb | NH2 | low | 35 | 0.264415 | CC | 2 | AA | -- | TT | CC | TT |
| P7RCC8 | NH2_Barb | NH2 | low | 33 | 0.2554 | CC | 2 | TT | -- | GG | CC | TT |
| P7RDC3 | NH2_Barb | NH2 | high | 45 | 0.26756 | CC | 2 | AA | T- | GG | NA | NA |
| P7RDC5 | NH2_Barb | NH2 | low | 36 | 0.2540675 | CC | 2 | AA | T- | GG | NA | NA |
| P7REC1 | NH2_Barb | NH2 | high | 43 | 0.2509475 | CC | 2 | AA | TT | GG | CC | TT |
| P7REC10 | NH2_Barb | NH2 | low | 34 | 0.266085 | AA | 0 | AA | TT | TT | TT | CC |
| P7REC5 | NH2_Barb | NH2 | high | 44 | 0.2623275 | CA | 1 | AA | T- | TT | TT | CC |
| P7REC6 | NH2_Barb | NH2 | high | 44 | 0.2675825 | CC | 2 | AA | TT | GG | CC | TT |
| P7REC7 | NH2_Barb | NH2 | low | 34 | 0.2631 | AA | 0 | AA | TT | TT | TT | CC |
| P7RGC3 | NH2_Barb | NH2 | high | 44 | 0.2759375 | CA | 1 | AA | TT | GT | TT | CC |

**Table S7.**
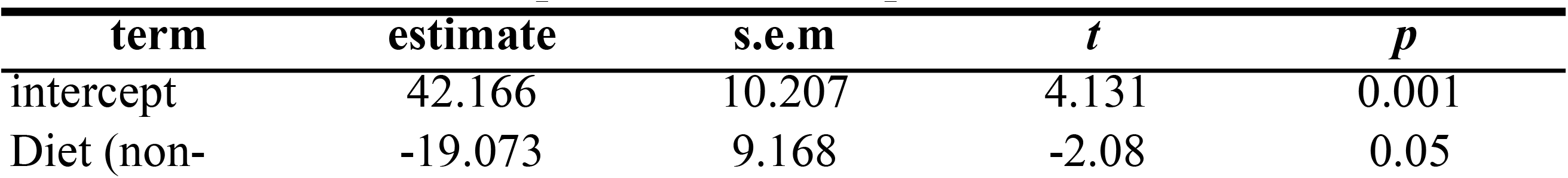
Summary of the phylogenetic least squares regression model testing for the effects of larval diet (herbivorous and non-herbivorous) on ovipositor bristle number per valve. For larval diet categorization, the reference category is ‘herbivorous.’ This analysis only includes the species reported in Table S2 that have both larval diet and ovipositor bristle counts available. F1, 21= 4.328, p=0.049, n= 23 species, λ=1.018.

| **term** | **estimate** | **s.e.m** | ***t*** | ***p*** |
| --- | --- | --- | --- | --- |
| intercept | 42.166 | 10.207 | 4.131 | 0.001 |
| Diet (non- | -19.073 | 9.168 | -2.08 | 0.05 |

**Table S8.**
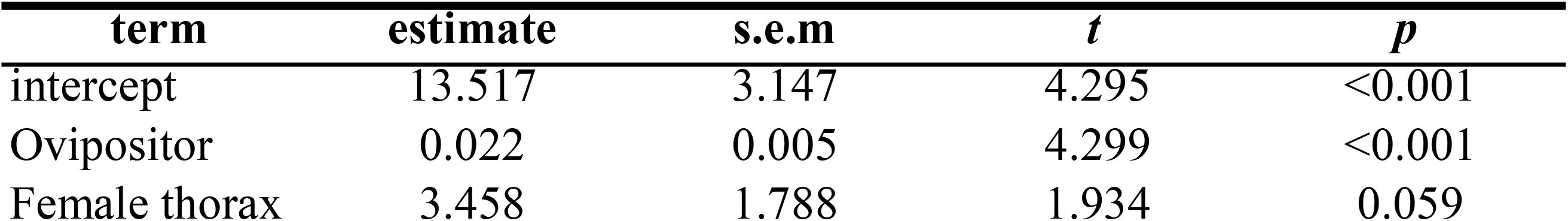
Summary of the regression model testing for the effects of ovipositor length and female thorax length on ovipositor bristle number. Data for 67 species were taken from Craddock et al. 2018, with additional data obtained for 4 species in this study. All species and measurements are listed in Table S3. Bristle counts were averaged across individuals within a species and across both ovipositor valves, when present. R2 =0.488, F2, 53=25.24, p<0.001.

| **term** | **estimate** | **s.e.m** | ***t*** | ***p*** |
| --- | --- | --- | --- | --- |
| intercept | 13.517 | 3.147 | 4.295 | <0.001 |
| Ovipositor | 0.022 | 0.005 | 4.299 | <0.001 |
| Female thorax | 3.458 | 1.788 | 1.934 | 0.059 |

**Table S9.**
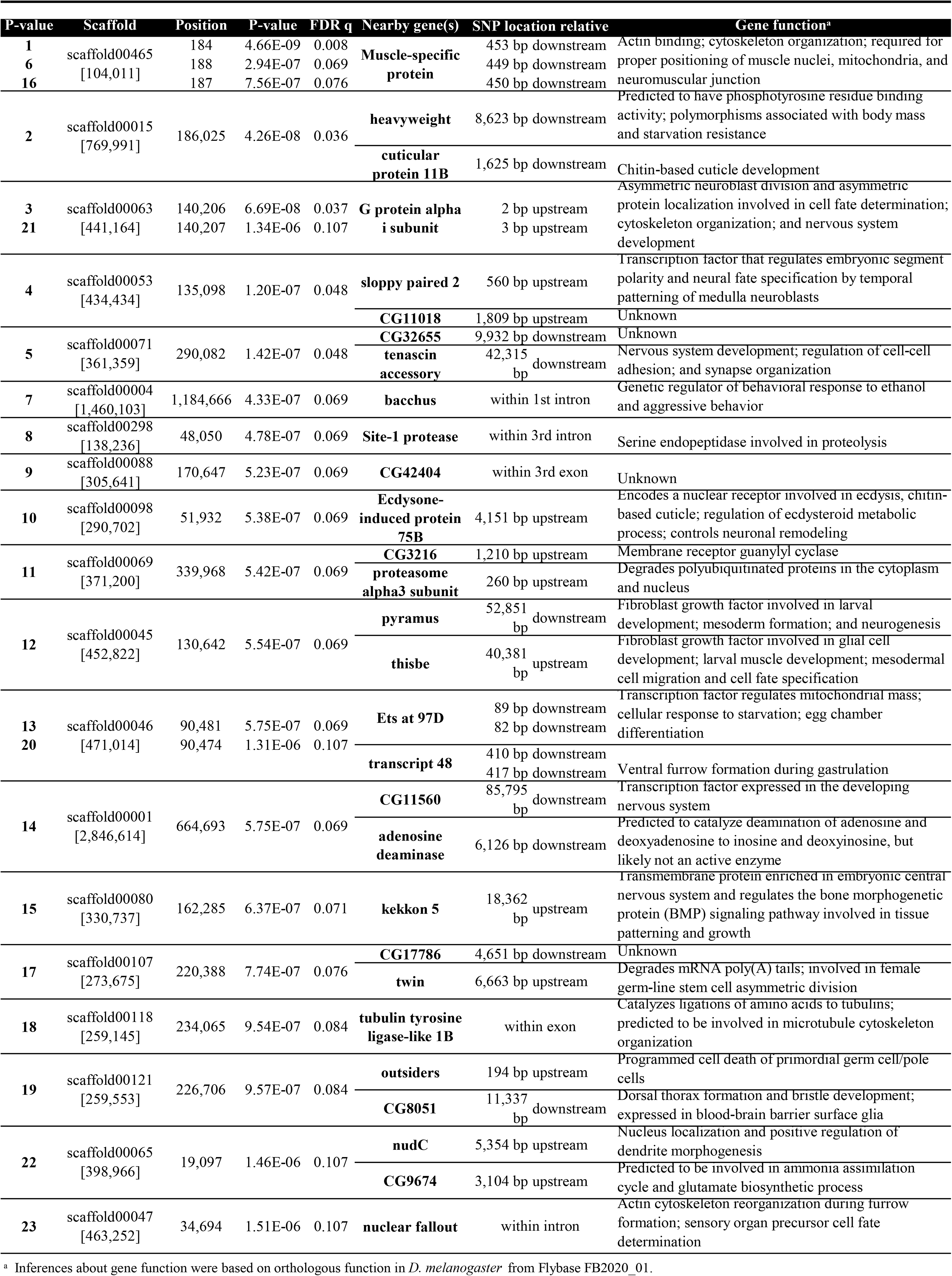
Top SNPs associated with variation in ovipositor bristle number from the pool genome-wide association study are shown in descending P-value ranking.

**Table S10.**
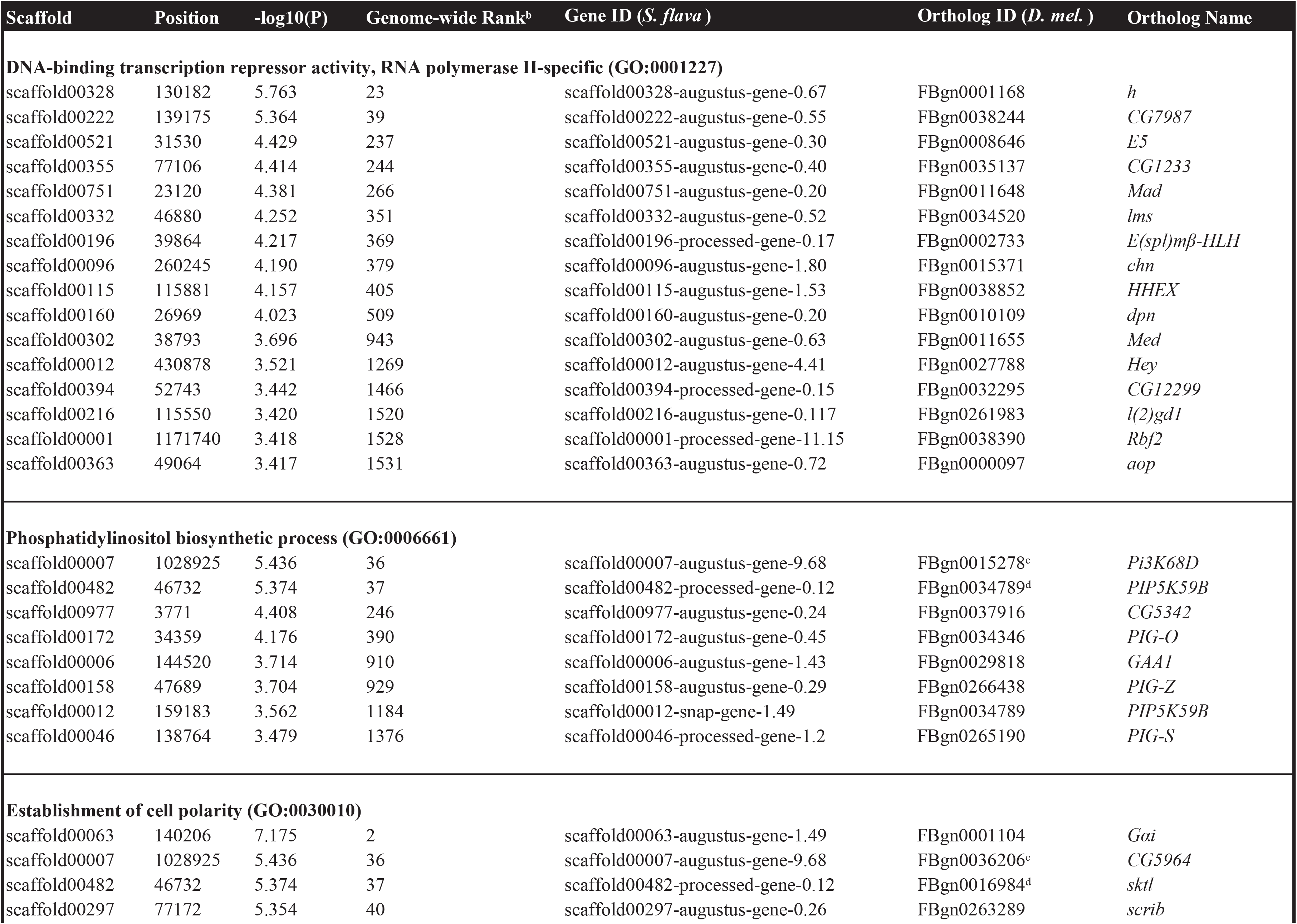
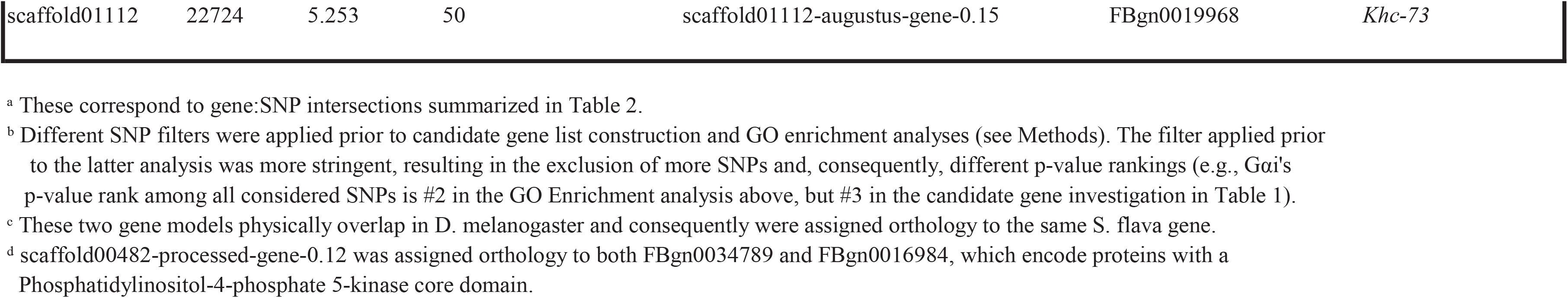
Genes annotated with enriched gene ontology (GO) terms that intersected the most significant pool-GWAS SNPs.a

**Table S11.** Summary of the regression model testing for the additive effects of a candidate SNP on ovipositor bristle number from individually sequenced S. flava flies. The SNP was identified from the pool-GWAS on ovipositor bristle number, and is located upstream of the neural development gene G alpha i subunit (Gαi). The model also tests for the effects of peg pool (high or low), ovipositor length, population/host plant (which could not be disentangled; NH1/Barbarea vulgaris/Turritis glabra, NH2/B. vulgaris, or NH2/T. glabra. For peg pool categorization, the reference category is ‘high,’ and for host plant/population, it is ‘NH1/Barbarea vulgaris/Turritis glabra’. Sequence data for this analysis has been deposited on Genbank (accession no. MH884655-MH884734). R2 =0.919, F5, 68=154.9, p<0.001, n= 74 individuals.

| term | estimate | s.e.m | *t* | *p* |
| --- | --- | --- | --- | --- |
| intercept | 29.983 | 3.446 | 8.702 | <0.001 |
| SNP | 0.583 | 0.203 | 2.876 | 0.005 |
| Peg pool (low) | -8.11 | 0.325 | -24.949 | <0.001 |
| Ovipositor length | 50.972 | 13.748 | 3.708 | <0.001 |
| Population/Host plant (NH2/*B.* | -0.889 | 0.392 | -2.27 | 0.026 |
| Population/Host plant (NH2/*T. glabra*) | 0.556 | 0.391 | 1.422 | 0.16 |

**Table S12.** Summary of the regression model testing for the effects of a candidate SNP (additive), peg pool (high or low), and population/host plant (NH1/Barbarea vulgaris/Turritis glabra, NH2/B. vulgaris, or NH2/T. glabra) on ovipositor length from individually sequenced S. flava flies. The candidate SNP was identified from the pool- GWAS on ovipositor bristle number, and is located upstream of the neural development gene G alpha i subunit (Gαi). For peg pool categorization, the reference category is ‘high,’ and for host plant/population, it is ‘NH1/Barbarea vulgaris/Turritis glabra’. Sequence data for this analysis has been deposited on Genbank (accession no. MH884655-MH884734). R2 =0.108, F4, 69=2.085, p=0.092, n= 74 species.

| **term** | **estimate** | **s.e.m** | ***t*** | ***p*** |
| --- | --- | --- | --- | --- |
| intercept | 0.249 | 0.003 | 74.431 | <0.001 |
| SNP | <0.001 | 0.002 | 0.177 | 0.86 |
| Peg pool (low) | <0.001 | 0.003 | 0.013 | 0.99 |
| Population/Host plant (NH2/B. vulgaris) | 0.009 | 0.003 | 2.81 | 0.006 |
| Population/Host plant (NH2/T. glabra) | 0.003 | 0.003 | 0.956 | 0.342 |

**Table S13.** Linkage disequilibrium estimates between the focal SNP identified from pool-GWA (locus 214), located upstream of the neural development gene Gai, and neighboring variants identified from individual re-sequencing. Delta values estimate correlations among unphased alleles. Strong LD values (| δ | > 0.5 (p value < 0.01) are shown in bold italics.

| Variant locus | Delta $\delta$ | | | | T2 | df | P-value |
| --- | --- | --- | --- | --- | --- | --- | --- |
|  | AA | Aa | aA | aa |  |  |  |
| 49 | 0.1559533 | ***0.560263*** | ***-0.8721695*** | 0.1559533 | 230.27 | 1 | 5.20E-52 |
| 213 | -0.229821 | ***0.6081994*** | -0.1485573 | -0.229821 | 106.03 | 1 | 7.26E-25 |
| 215 | -0.2529218 | 0.2529218 | 0.2529218 | -0.2529218 | 48.64 | 1 | 3.08E-12 |
| 235 | -0.2907232 | 0.2907232 | 0.2907232 | -0.2907232 | 65.01 | 1 | 7.47E-16 |
| 236 | 0.2849708 | -0.2849708 | -0.2849708 | 0.2849708 | 68.1 | 1 | 1.55E-16 |

## References

1. Mitter C, Farrell B, Wiegmann B. 1988 The Phylogenetic Study of Adaptive Zones: Has Phytophagy Promoted Insect Diversification? Am. Nat. 132, 107–128.

2. Wiens JJ, Lapoint RT, Whiteman NK. 2015 Herbivory increases diversification across insect clades. Nat. Commun. 6, 8370.

3. Schoonhoven LM, Van Loon B, van Loon JJA, Dicke M. 2005 Insect-Plant Biology. OUP Oxford.

4. Southwood TRE. 1972 insect/plant relationship--an evolutionary perspective. Roy Entomol Soc London Symp 1972, 6.

5. Wiegmann BM et al. 2011 Episodic radiations in the fly tree of life. Proc. Natl. Acad. Sci. U. S. A. 108, 5690–5695.

6. Marazzi B, Ané C, Simon MF, Delgado-Salinas A, Luckow M, Sanderson MJ. 2012 Locating evolutionary precursors on a phylogenetic tree. Evolution 66, 3918–3930.

7. Rabosky DL. 2017 Phylogenetic tests for evolutionary innovation: the problematic link between key innovations and exceptional diversification. Philos. Trans. R. Soc. Lond. B Biol. Sci. 372. (doi:10.1098/rstb.2016.0417)

8. Eiseman C, Charney N, Carlson J. 2010 Tracks & Sign of Insects & Other Invertebrates: A Guide to North American Species. Stackpole Books.

9. Connor EF, Taverner MP. 1997 The Evolution and Adaptive Significance of the Leaf- Mining Habit. Oikos 79, 6–25.

10. Atallah J, Teixeira L, Salazar R, Zaragoza G, Kopp A. 2014 The making of a pest: the evolution of a fruit-penetrating ovipositor in Drosophila suzukii and related species. Proc. Biol. Sci. 281, 20132840.

11. Gloss AD et al. 2019 Evolution of herbivory remodels a Drosophila genome. bioRxiv., 767160. (doi:10.1101/767160)

12. Thomson JA, Jackson MJ, Bock IR. 1982 Contrasting resource utilisation in two Australian species of Drosophila Fallen (Diptera) feeding on the bracken fern Pteridium scopoli. Aust. J. Entomol. 21, 29–30.

13. Yassin A, Debat V, Bastide H, Gidaszewski N, David JR, Pool JE. 2016 Recurrent specialization on a toxic fruit in an island Drosophila population. Proc. Natl. Acad. Sci. U. S. A. 113, 4771–4776.

14. Groen SC, Whiteman NK. 2016 Using Drosophila to study the evolution of herbivory and diet specialization. Curr Opin Insect Sci 14, 66–72.

15. Kim BY et al. 2021 Highly contiguous assemblies of 101 drosophilid genomes. Elife 10. (doi:10.7554/eLife.66405)

16. Shakeel M, He XZ, Martin NA, Hanan A, Wang Q. 2009 Diurnal periodicity of adult eclosion mating and oviposition of the european leafminer Scaptomyza flava (Falln)(Diptera Drosophilidae). N. Z. Plant Prot. 62, 80–85.

17. Martin N. 2014 Scaptomyza (Bunostoma) flavella (Diptera: Drosophilidae) and the evolution of leaf mining. 1 47, 8–11.

18. Norga KK et al. 2003 Quantitative analysis of bristle number in Drosophila mutants identifies genes involved in neural development. Curr. Biol. 13, 1388–1396.

19. Sham P, Bader JS, Craig I, O’Donovan M, Owen M. 2002 DNA Pooling: a tool for large- scale association studies. Nat. Rev. Genet. 3, 862–871.

20. Bastide H, Betancourt A, Nolte V, Tobler R, Stöbe P, Futschik A, Schlötterer C. 2013 A genome-wide, fine-scale map of natural pigmentation variation in Drosophila melanogaster. PLoS Genet. 9, e1003534.

21. Katoh T, Izumitani HF, Yamashita S, Watada M. 2017 Multiple origins of Hawaiian drosophilids: phylogeography of Scaptomyza hardy (Diptera: Drosophilidae). Entomol. Sci. 20, 33–44.

22. Gloss AD et al. 2014 Evolution in an ancient detoxification pathway is coupled with a transition to herbivory in the drosophilidae. Mol. Biol. Evol. 31, 2441–2456.

23. Lapoint RT, O’Grady PM, Whiteman NK. 2013 Diversification and dispersal of the Hawaiian Drosophilidae: the evolution of Scaptomyza. Mol. Phylogenet. Evol. 69, 95–108.

24. Stamatakis A. 2006 RAxML-VI-HPC: maximum likelihood-based phylogenetic analyses with thousands of taxa and mixed models. Bioinformatics 22, 2688–2690.

25. Ronquist F, Huelsenbeck JP. 2003 MrBayes 3: Bayesian phylogenetic inference under mixed models. Bioinformatics 19, 1572–1574.

26. Bouckaert R, Heled J, Kühnert D, Vaughan T, Wu C-H, Xie D, Suchard MA, Rambaut A, Drummond AJ. 2014 BEAST 2: a software platform for Bayesian evolutionary analysis. PLoS Comput. Biol. 10, e1003537.

27. Butler MA, King AA. 2004 Phylogenetic Comparative Analysis: A Modeling Approach for Adaptive Evolution. Am. Nat. 164, 683–695.

28. Paradis E, Claude J, Strimmer K. 2004 APE: Analyses of Phylogenetics and Evolution in R language. Bioinformatics. 20, 289–290. (doi:10.1093/bioinformatics/btg412)

29. Kembel SW, Cowan PD, Helmus MR, Cornwell WK, Morlon H, Ackerly DD, Blomberg SP, Webb CO. 2010 Picante: R tools for integrating phylogenies and ecology. Bioinformatics 26, 1463–1464.

30. Harmon LJ, Weir JT, Brock CD, Glor RE, Challenger W. 2008 GEIGER: investigating evolutionary radiations. Bioinformatics 24, 129–131.

31. Pagel M. 1999 Inferring the historical patterns of biological evolution. Nature 401, 877–884.

32. Revell LJ. 2012 phytools: an R package for phylogenetic comparative biology (and other things). Methods Ecol. Evol. 3, 217–223.

33. Craddock EM, Kambysellis MP, Franchi L, Francisco P, Grey M, Hutchinson A, Nanhoo S, Antar S. 2018 Ultrastructural variation and adaptive evolution of the ovipositor in the endemic Hawaiian Drosophilidae. J. Morphol. 279, 1725–1752.

34. Schlötterer C, Tobler R, Kofler R, Nolte V. 2014 Sequencing pools of individuals—mining genome-wide polymorphism data without big funding. Nat. Rev. Genet. 15, 749–763.

35. Kofler R, Pandey RV, Schlötterer C. 2011 PoPoolation2: identifying differentiation between populations using sequencing of pooled DNA samples (Pool-Seq). Bioinformatics 27, 3435– 3436.

36. Thurmond J et al. 2019 FlyBase 2.0: the next generation. Nucleic Acids Res. 47, D759– D765.

37. Kofler R, Schlötterer C. 2012 Gowinda: unbiased analysis of gene set enrichment for genome-wide association studies. Bioinformatics 28, 2084–2085.

38. Schaefer M, Petronczki M, Dorner D, Forte M, Knoblich JA. 2001 Heterotrimeric G proteins direct two modes of asymmetric cell division in the Drosophila nervous system. Cell 107, 183–194.

39. Schaid DJ. 2004 Evaluating associations of haplotypes with traits. Genet. Epidemiol. 27, 348–364.

40. Reynolds N, O’Shaughnessy A, Hendrich B. 2013 Transcriptional repressors: multifaceted regulators of gene expression. Development 140, 505–512.

41. Usui K, Goldstone C, Gibert J-M, Simpson P. 2008 Redundant mechanisms mediate bristle patterning on the Drosophila thorax. Proc. Natl. Acad. Sci. U. S. A. 105, 20112–20117.

42. Balakrishnan SS, Basu U, Raghu P. 2015 Phosphoinositide signalling in Drosophila. Biochim. Biophys. Acta 1851, 770–784.

43. Janardan V, Sharma S, Basu U, Raghu P. 2020 A Genetic Screen in Drosophila To Identify Novel Regulation of Cell Growth by Phosphoinositide Signaling. G3 **10**, 57–67.

44. Hassan BA, Prokopenko SN, Breuer S, Zhang B, Paululat A, Bellen HJ. 1998 skittles, a Drosophila phosphatidylinositol 4-phosphate 5-kinase, is required for cell viability, germline development and bristle morphology, but not for neurotransmitter release. Genetics 150, 1527–1537.

45. Schweisguth F. 2015 Asymmetric cell division in the Drosophila bristle lineage: from the polarization of sensory organ precursor cells to Notch-mediated binary fate decision. Wiley Interdisciplinary Reviews: Developmental Biology. 4, 299–309. (doi:10.1002/wdev.175)

46. Green JE, Cavey M, Médina Caturegli E, Aigouy B, Gompel N, Prud’homme B. 2019 Evolution of Ovipositor Length in Drosophila suzukii Is Driven by Enhanced Cell Size Expansion and Anisotropic Tissue Reorganization. Curr. Biol. 29, 2075–2082.e6.

47. Stern DL, Orgogozo V. 2008 The loci of evolution: how predictable is genetic evolution? Evolution 62, 2155–2177.

48. Bhat KM, van Beers EH, Bhat P. 2000 Sloppy paired acts as the downstream target of wingless in the Drosophila CNS and interaction between sloppy paired and gooseberry inhibits sloppy paired during neurogenesis. Development 127, 655–665.

49. Orgogozo V, Schweisguth F, Bellaïche Y. 2001 Lineage, cell polarity and inscuteable function in the peripheral nervous system of the Drosophila embryo. Development 128, 631– 643.

50. Taylor BJ. 1989 Sexually dimorphic neurons in the terminalia of Drosophila melanogaster: *I.* Development of sensory neurons in the genital disc during metamorphosis. J. Neurogenet. 5, 173–192.

51. Nagy O et al. 2018 Correlated Evolution of Two Copulatory Organs via a Single cis- Regulatory Nucleotide Change. Curr. Biol. 28, 3450–3457.e13.

52. Marcellini S, Simpson P. 2006 Two or four bristles: functional evolution of an enhancer of scute in Drosophilidae. PLoS Biol. 4, e386.

53. Robin C, Lyman RF, Long AD, Langley CH, Mackay TFC. 2002 hairy: A quantitative trait locus for drosophila sensory bristle number. Genetics 162, 155–164.

54. Hagen JFD et al. 2021 Unraveling the Genetic Basis for the Rapid Diversification of Male Genitalia between Drosophila Species. Mol. Biol. Evol. 38, 437–448.

55. Lamichhaney S, Han F, Berglund J, Wang C, Almén MS, Webster MT, Grant BR, Grant PR, Andersson L. 2016 A beak size locus in Darwin’s finches facilitated character displacement during a drought. Science 352, 470–474.

56. Linnen CR, Kingsley EP, Jensen JD, Hoekstra HE. 2009 On the origin and spread of an adaptive allele in deer mice. Science 325, 1095–1098.

57. Rost S et al. 2004 Mutations in VKORC1 cause warfarin resistance and multiple coagulation factor deficiency type 2. Nature 427, 537–541.

## References

1. Nylander JAA, Ronquist F, Huelsenbeck JP, Nieves-Aldrey JL. 2004 Bayesian phylogenetic analysis of combined data. Syst. Biol. 53, 47–67.

2. Swofford DL, Sullivan J. 2003 Phylogeny inference based on parsimony and other methods using PAUP*. The phylogenetic handbook: a practical approach to DNA and protein phylogeny 7, 160–206.

3. Katoh T, Izumitani HF, Yamashita S, Watada M. 2017 Multiple origins of Hawaiian drosophilids: phylogeography of Scaptomyza hardy (Diptera: Drosophilidae). Entomol. Sci. 20, 33–44.

4. Bouckaert R, Heled J, Kühnert D, Vaughan T, Wu C-H, Xie D, Suchard MA, Rambaut A, Drummond AJ. 2014 BEAST 2: a software platform for Bayesian evolutionary analysis. PLoS Comput. Biol. 10, e1003537.

5. Ayres DL et al. 2012 BEAGLE: an application programming interface and high- performance computing library for statistical phylogenetics. Syst. Biol. 61, 170–173.

6. Rambaut A, Suchard MA, Xie D, Drummond AJ. 2014 Tracer v1. 6. Computer program and documentation distributed by the author.

7. Lynch M, Walsh B, Others. 1998 Genetics and analysis of quantitative traits. Sinauer Sunderland, MA.

8. Bolger AM, Lohse M, Usadel B. 2014 Trimmomatic: a flexible trimmer for Illumina sequence data. Bioinformatics 30, 2114–2120.

9. Li H, Durbin R. 2009 Fast and accurate short read alignment with Burrows–Wheeler transform. Bioinformatics 25, 1754–1760.

10. Schlötterer C, Tobler R, Kofler R, Nolte V. 2014 Sequencing pools of individuals—mining genome-wide polymorphism data without big funding. Nat. Rev. Genet. 15, 749–763.

11. Li H et al. 2009 The Sequence Alignment/Map format and SAMtools. Bioinformatics 25, 2078–2079.

12. Kofler R, Pandey RV, Schlötterer C. 2011 PoPoolation2: identifying differentiation between populations using sequencing of pooled DNA samples (Pool-Seq). Bioinformatics 27, 3435– 3436.

13. Yang J et al. 2011 Genomic inflation factors under polygenic inheritance. Eur. J. Hum. Genet. 19, 807–812.

14. Devlin B, Roeder K. 1999 Genomic control for association studies. Biometrics 55, 997– 1004.

15. Thoen MPM et al. 2017 Genetic architecture of plant stress resistance: multi-trait genome- wide association mapping. New Phytol. 213, 1346–1362.

16. Lunter G, Goodson M. 2011 Stampy: a statistical algorithm for sensitive and fast mapping of Illumina sequence reads. Genome Res. 21, 936–939.

17. McKenna A et al. 2010 The Genome Analysis Toolkit: a MapReduce framework for analyzing next-generation DNA sequencing data. Genome Res. 20, 1297–1303.

18. Quinlan AR, Hall IM. 2010 BEDTools: a flexible suite of utilities for comparing genomic features. Bioinformatics 26, 841–842.

19. Feder AF, Petrov DA, Bergland AO. 2012 LDx: estimation of linkage disequilibrium from high-throughput pooled resequencing data. PLoS One 7, e48588.

20. Li L, Stoeckert CJ Jr, Roos DS. 2003 OrthoMCL: identification of ortholog groups for eukaryotic genomes. Genome Res. 13, 2178–2189.

21. Drosophila 12 Genomes Consortium et al. 2007 Evolution of genes and genomes on the Drosophila phylogeny. Nature 450, 203–218.

22. Carlson M, Falcon S, Pages H, Li N. 2017 GO. db: A set of annotation maps describing the entire Gene Ontology. R package version 3, 10–18129.

23. Gentleman RC et al. 2004 Bioconductor: open software development for computational biology and bioinformatics. Genome Biol. 5, R80.

24. Kofler R, Schlötterer C. 2012 Gowinda: unbiased analysis of gene set enrichment for genome-wide association studies. Bioinformatics 28, 2084–2085.

25. Kvon EZ, Kazmar T, Stampfel G, Yáñez-Cuna JO, Pagani M, Schernhuber K, Dickson BJ, Stark A. 2014 Genome-scale functional characterization of Drosophila developmental enhancers in vivo. Nature 512, 91–95.

26. Dmitriev DA, Rakitov RA. 2008 Decoding of superimposed traces produced by direct sequencing of heterozygous indels. PLoS Comput. Biol. 4, e1000113.

